# Multi-organ single-cell transcriptomics of immune cells uncovered organ-specific gene expression and functions

**DOI:** 10.1101/2023.05.29.542692

**Authors:** Maria Tsagiopoulou, Sonal Rashmi, Sergio Aguilar, Juan Nieto, Ivo G. Gut

**Affiliations:** CNAG-CRG, Centre for Genomic Regulation (CRG), Barcelona Institute of Science and Technology (BIST), Barcelona, Spain; Universitat Pompeu Fabra (UPF), Barcelona, Spain

## Abstract

Despite the large number of publicly available single-cell datasets, there is a limited understanding of the distinct resident immune cells and their concomitant features in diverse human organs. To address this, we compiled a dataset of 114,275 *CD45+* immune cells from 14 organs from healthy donors. Although the transcriptome of immune cells is constant across organs, organ-specific gene expression changes were detected revealing unique expression in certain organs (*GTPX3* in kidney, *DNTT* and *ACVR2B* in thymus). These alterations are associated with different transcriptional factor activities and pathways including metabolism. *TNF-α* signaling through the *NFkB* pathway was found in various organs and immune compartments including distinct expression profiles of *NFkB* family genes and their target genes such as cytokines indicating their role in cell positioning. Taken together, immune cells not only protect the organs but also adapt to the host organ environment and contribute to its function and homeostasis.

## Introduction

Immune response is classified into: (i) innate (dendritic, macrophages, granulocytes, NK cells) and (ii) adaptive (B and T cells). Most immune cells are produced in the bone marrow and reach other organs through blood. Knowledge of human immune cells is derived from peripheral blood and there is limited understanding of the diversity, specificity and behavior of immune cell across multiple organs. State-of-the-art technologies to analyze individual cells such as single-cell RNA-seq (scRNA-seq) have been used to study organ-specific immunity^1^, ^2^.

Public datasets of scRNA-seq are increasing through international collaborative efforts such as the Human Cell Atlas (HCA) charting the cell types in the healthy body, across time from fetal development to adulthood, and eventually to old age^3^. There has been little attempt to integrate the scRNA-seq data from different resources^4^. However, recent publications managed to map the cell types across different human tissues and report pan-tissue atlases from a different perspective^5^, ^6^, ^7^, ^8^. Dominguez Conde et al. focused on the immune cell analysis in different tissues and characterized in depth the cells concluding to a hundred immune populations^5^ and focusing mainly on macrophages and memory T cells. Large-scale multi-organ meta-atlases on the specific or shared genes and functions of the immune cells regarding their host organs have not been reported.

As the first attempts to characterize the immunity of human organs are starting, many publicly available data could be integrated. In this context, we analyzed the transcriptional landscape of 114,275 immune cells in 14 different organs addressing the unique and shared features characterizing the different organs. In this study, we aimed to investigate whether immune cells undergo any changes while residing in the host organ, if they utilize any components of the host organ, and whether they play a role in the host organ beyond their defensive function. Specifically, we sought to understand the mechanisms underlying immune cells’ entry into host organs. We report gene expression changes of certain immune cells across various organs with certain genes exhibiting high levels of specificity to particular organs. The altered genes were associated with specific or shared transcription factor activity and biological pathways, such as the *NFkB* signaling pathway, which influences cytokines expression.

## Results

### Multi-organs immune cells map distinct immune cell populations

We aggregated 162 healthy donor samples from 14 organs (**Supplementary Table 1)** and 12 different projects (1/12 from liver transplantation) from HCA portal focusing on *CD45*+ compartment (**Fig. 1a, Supplementary-Fig. 1a-c**). The dataset after the quality control process comprised 114,275 goodquality *CD45*+ cells. Notably, the *CD45*+ selection led to a reduction of the progenitor lineage due to low expression of *CD45 (***Supplementary-Fig. c**).

**Fig. 1.**
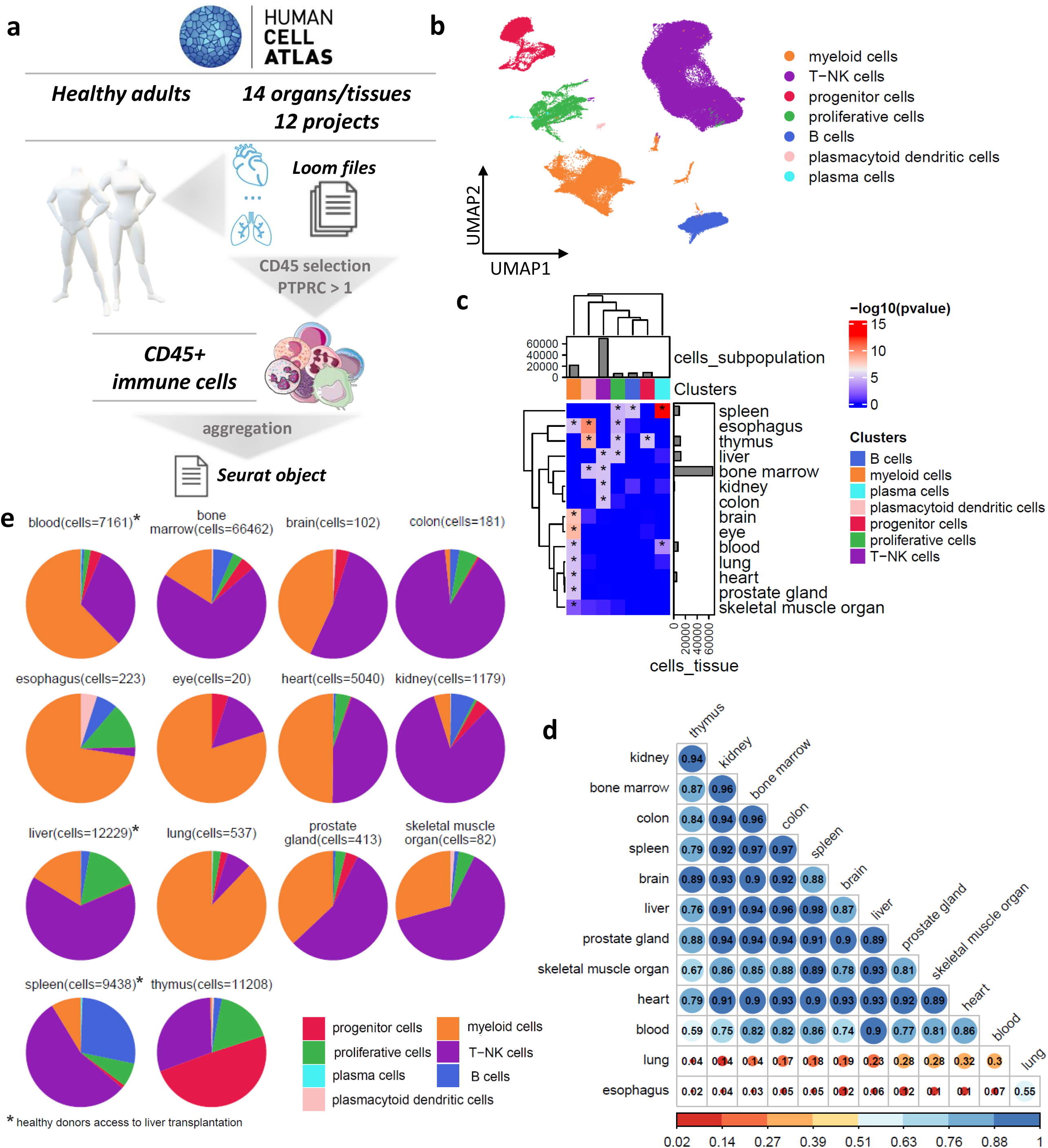
Single-cell transcriptomics of immune cells from different organs reveal the common immune cell subpopulation. a, Graphical representation of the study. **b**, The UMAP plot of the immune cells. **c**, Heatmap showing the -log10(p-value) of the hypergeometric distribution between the immune cell subpopulations and the different organs. The asterisks highlight the statistically significant enrichment (p-value<0.05). **e**, Pie charts displaying the frequencies of the immune cell subtypes across the different organs. **d**, Pearson correlation plot across the different organs.

We aim to understand the immune cell composition heterogeneity in different organs (prostate gland, eye, heart, skeletal muscle organ, blood, liver, brain, kidney, colon, esophagus, and lung) including lymphoid organs (thymus, bone marrow, spleen). We identified the major cell types: T-NK (n=69,285), myeloid (n=21,451), B (n=7,342), plasmacytoid dendritic (n=349) and plasma (n=162) cells and two states that thread across cell types: progenitors (n=9,422) and proliferative cells (n=6,912) (**Fig. 1b, Supplementary-Fig. 1d-f)**. The over-presenting organ in terms number of immune cells was the bone marrow (n=66,465) followed by the liver (n=12,513), thymus (n=11,208), and spleen (n=9,743) (**Table 1**). Brain, eye, blood, lung, prostate gland, heart, esophagus, and skeletal muscle organ showed statistically significant enrichment (hypergeometric distribution, p<0.05) of myeloid cells (**Fig. 1c-e**). Spleen was associated with B cells and together with blood was associated with plasma cells. The spleen, liver, esophagus and thymus were enriched in proliferative cells. Bone marrow, liver, kidney and colon were enriched in T cells and bone marrow showed additional enrichment in plasmacytoid dendritic cells (DC). Then, we compared immune cells expression profiles from different organs using the Pearson correlation coefficient (r) (**Fig. 1d**). The results (Pearson’s-r: minimum 0.02-maximum 0.96) showed completely uncorrelated profiles of lung and esophagus compared to the remaining organs with a range of Pearson’s-r 0.04-0.3 and 0.02-0.12 respectively, however a weak correlation between them was found (Pearson’s-r= 0.55). The best correlation was between bone marrow and the colon (Pearson’s-r= 0.96) and kidney (Pearson’s-r= 0.96). A public annotation of the tonsil was used to characterize the immune cells in detail^9^. B and myeloid cells had the same cell-annotation, but the T-NK cells were more finely characterized using this reference (**Supplementary-Fig. 2**).

**Table 1.**
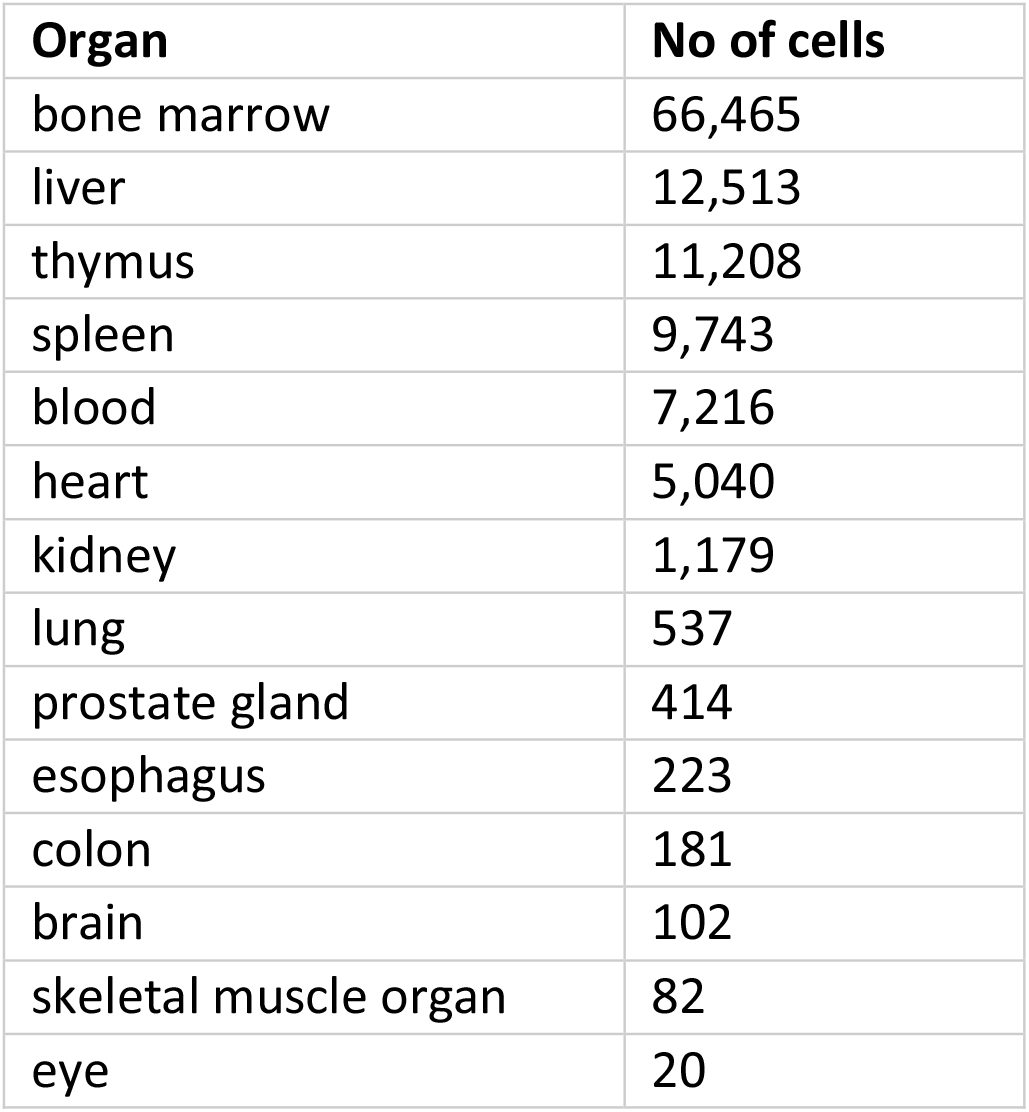
The studied organs and the number of CD45+ immune cells.

### B and progenitor (lymphoid and myeloid lineages) cells analysis revealed thymus properties, thymic B cells and an association with kidney

The 7,342 B cells (6.5% of detected immune cells) grouped into five clusters, including one cluster of immature stage expressing *SOX4* (pre B cells) (**Fig. 2a, Supplementary-Fig. 3a**). Mature memory B cells clusters characterized by *CD27*+ B cells (n=2,550) combined with memory B cells (n=1,819) were the over-presented subpopulations with 59.5% of the defining B cells (**Fig. 2b**). Of the 7,342 B cells 3,963 were found in bone marrow, 2,610 in the spleen, 316 in the liver and interestingly 258 in the thymus. A pre-B cell (n=312) cluster was organ-specific since the bone marrow plays the main role in the production of B cells (**Fig. 2b, Supplementary-Fig. 3b**). Also, bone marrow B cells were statistically enriched (hypergeometric distribution, p<0.001) in naive B cells (n=1,604). Moreover, the thymic B cells were enriched mainly in mature memory B cells and memory B cells highlighting their critical role in the maturation of T cells^10^. The best correlations were observed between spleen and liver, liver and blood, thymus and kidney (Pearson’s-r: 0.99) (**Supplementary-Fig. 3c**) while the lowest correlation was between heart, kidney and thymus with liver and spleen. Focused analysis of mature and memory B cells from thymus showed the best correlation with the cells located in bone marrow (**Supplementary-Fig. 3d**) supporting evidence of thymic B cells origin.

**Fig. 2.**
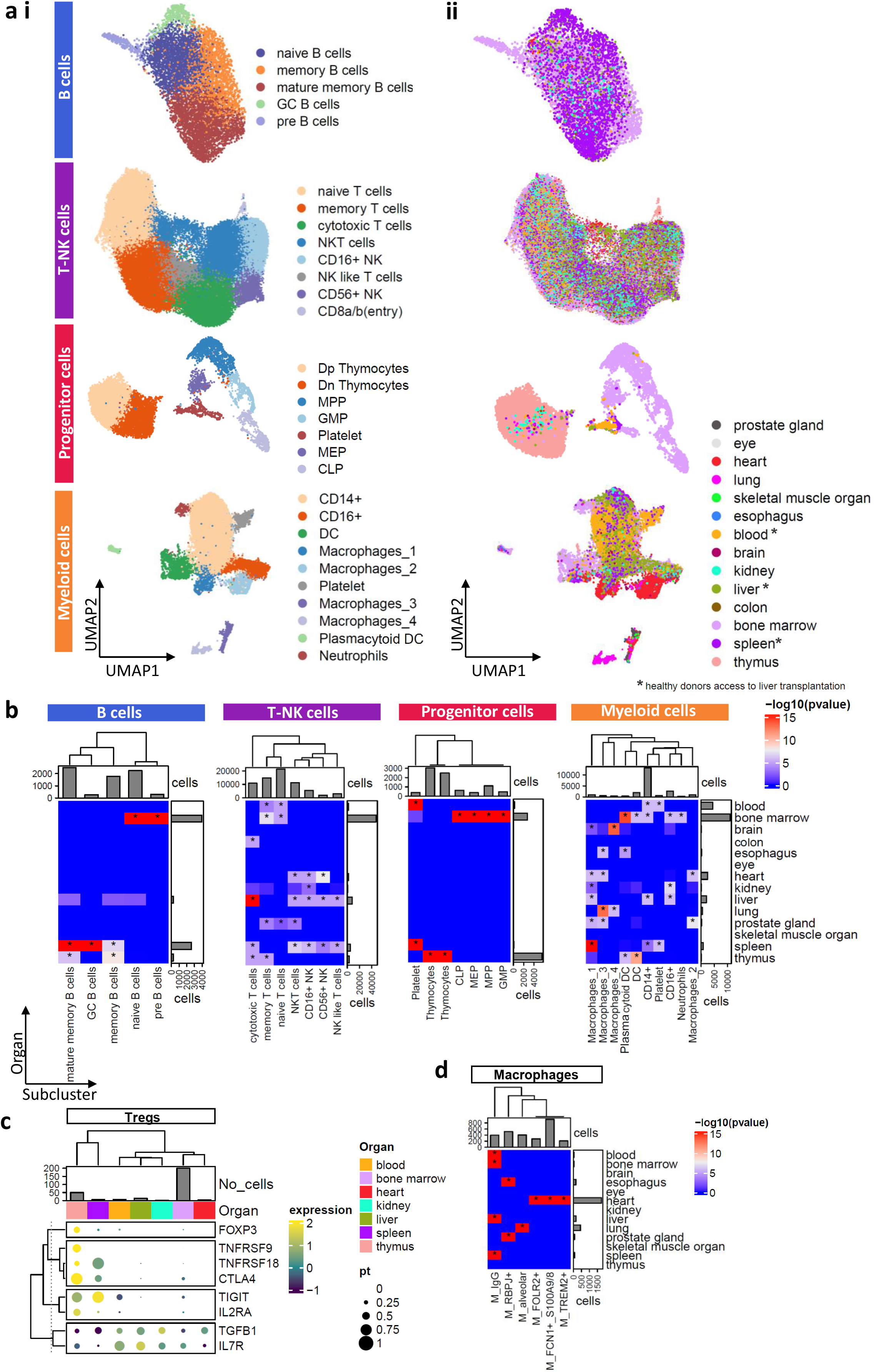
Mapping the different immune cell types across the different organs. **a,** The UMAP plots of the (**i**) different immune cell types in (**ii**) different organs. The asterisks on organs highlighting the liver transplantation. **b**, Heatmap showing the -log10(p-value) of hypergeometric distribution between the immune cell subpopulations and the different organs. The asterisks highlight the statistically significant enrichment (p-value<0.05) **c**, Hierarchical clustering based on expression levels and percentage of cells expressing the Tregs markers. The barplot showing the number of Tregs across organs **d**, Heatmap showing the -log10(p-value) of hypergeometric distribution across macrophages and different organs.

We obtained 9,422 progenitor cells characterized by the expression of classical markers such as *SOX4*. As expected, the over-presented organs in this immune subtype were the thymus (n=5,524) and bone marrow (n=2,791) (**Fig. 2a, Supplementary-Fig. 3e-f**). The thymus was enriched in Dp and Dn thymocytes and bone marrow in myeloid and lymphoid lineage (**Fig. 2b, Supplementary-Fig. 3e-f**). The detected platelet cells (n=422) were associated with blood and spleen. Interestingly, the progenitor cells derived from the kidney were clustered with the thymocytes (46/47 progenitor cells from kidney).

### High diversity of T and myeloid cells and organ specificity in macrophages

T and NK (T-NK) cells were the over-presented immune cell types (n=69,285, 60% of detected immune cells) and were divided into eight clusters (**Fig. 2a, Supplementary-Fig. 4a-b**). Most cells (52.2%) were found in the naïve (n=21,360) and memory (n=14,784) clusters expressing *SELL* and were separated by the differential expression of activated genes such as *FOS* (**Fig. 2b, Supplementary-Fig. 4a**). Memory T cells were divided into *CD4*+, *CD8*+ or *LEF1*+ clusters with the last one being associated with thymus (**Supplementary-Fig. 4c-d**). The *CD8*+ cells were found in spleen and liver and the *CD4*+ in thymus, kidney, heart, prostate gland and bone marrow (hypergeometric distribution, p<0.001). Additionally, a Treg subcluster (n=283) was found in the *CD4*+ subpopulation showing distinct transcriptomic signatures per organ. ?hymic Tregs (natural Tregs-nTregs) had the highest levels of *FOXP3* and *CTLA4* expression (**Supplementary-Fig. 4e-h**). Tregs from spleen showed similar expression patterns with nTregs expressing *CTLA4, TNFRSF18* and *TIGIT* (**Fig. 2c**).

Five clusters associated with cytotoxicity (*CD56*+ NK, *CD16*+ NK, NK-T cells, NK like T cells and cytotoxic T cells), captures 47.5% of the defined T-NK cells. A cluster was characterized as CD8a/b (entry) cells and was fully enriched in the thymus (225/227 cells) (**Fig. 2b, Supplementary-Fig. 4a-b**). Of the 69,285 T-NK cells, 46,733 were found in bone marrow, 7,949 in the liver, 5,197 in the spleen, 3,383 in the thymus, 2,253 in blood and 2,247 in the heart (**Fig. 2b**). Bone marrow and blood were enriched in naïve and memory T cells. Spleen and liver, on the other hand, were enriched in the five clusters associated with cytotoxicity (**Fig. 2b, Supplementary-Fig. 4a**). The remaining organs were mainly enriched in another cluster of cytotoxic T-NK cells. T-NK cells expression profiles across organs showed high variability (Pearson’s-r: min=0.77, max=0.99) (**Supplementary-Fig. 4**,**i**). The best correlation was observed between the liver and spleen (Pearson’s-r: 0.99) since they were enriched in cytotoxic clusters and the most distinct organ was skeletal muscle (Pearson’s-r: 0.77-0.94).

A total of 21,800 myeloid cells were grouped into ten clusters including four transcriptionally distinct macrophage subpopulations (**Fig. 2a, Supplementary-Fig. 5a-c**). Sixty percent of the cells were characterized as *CD14*+, 13.4% as macrophages, 11.8% as *CD16*+ and 8.7% as DCs. We captured platelets (n=625) with blood and spleen, plasmacytoid DCs (n=348) and neutrophils (n=312) associated with bone marrow (hypergeometric distribution, p-value<0.05). Macrophages were enriched in spleen, lung and brain (**Fig. 2b**). Half of the myeloid cells were derived from bone marrow (n=10,926) followed by the myeloid cells from blood (n=4,460) and liver (n=1,993) (**Fig. 2b**). High heterogeneity was observed across the different organs in myeloid cells (Pearson′s r: 0.07-0.97) (**Supplementary-Fig. 5c**). The most distinct organs were the lung (r: 0.2-0.59) and the esophagus (r: 0.07-0.53) and the highest correlation was found between the spleen and liver (r=0.99).

Macrophages (n=2,933) analysis revealed four clusters including the most distinct subpopulation of alveolars (**Supplementary-Fig. 5d-f**). The highest number of myeloid cells was found in heart (n=1,755), followed by lung (n=463) and liver (n=203). Using gene markers, we identified M1- and M2- associated clusters (**Supplementary-Fig. 5f**). The inflammatory *FCN1*+_*S100A9*/*8* macrophages (n=986) were the over-presented subpopulation and mostly in heart (**Supplementary-Fig. 5f-h**). The macrophages from the heart were also associated with *TREM2*+ macrophages (**Fig. 2d, Supplementary-Fig. 5g**). M2 polarization markers were found in *FOLR2*+ (n=329), RBPJ+ (n=527) and alveolar (n=410) macrophages. Interestingly, a macrophage subcluster expressing *IGHG* genes without the presence of *CD79A* expression was observed in cells derived from the liver, spleen, blood, and a few from the bone marrow (**Supplementary-Fig. 5h**). This entity was identified as organ-resident macrophages, distinct from monocyte-derived macrophages due to differential expression of IgG receptors^11^.

### Organ- and immune-related expression of genes by differential expression analysis across the different organs

To gain insight into the organ-specificity of the cells, we sought to identify specific gene signatures for immune cells found in different organs by applying a threshold of 40 immune cells. We carried out differential expression analysis of each organ against all the others in each immune cell type category and we screened for uniquely overexpressed genes in a hierarchical cluster analysis. The gene set revealed for each organ was characterized as its gene signature and these signatures showed organ-specificity since there was no difference in the expression levels in intra-subtype analysis (*i*.*e*. in-depth characterization of the immune compartments) (**Fig. 3a**). The same analysis was repeated considering the down-regulated genes per organ in each immune cell type. No gene showed a unique reduction of the expression in certain organ however downregulation of genes was observed (**Supplementary-Fig. 6a**).

**Fig. 3.**
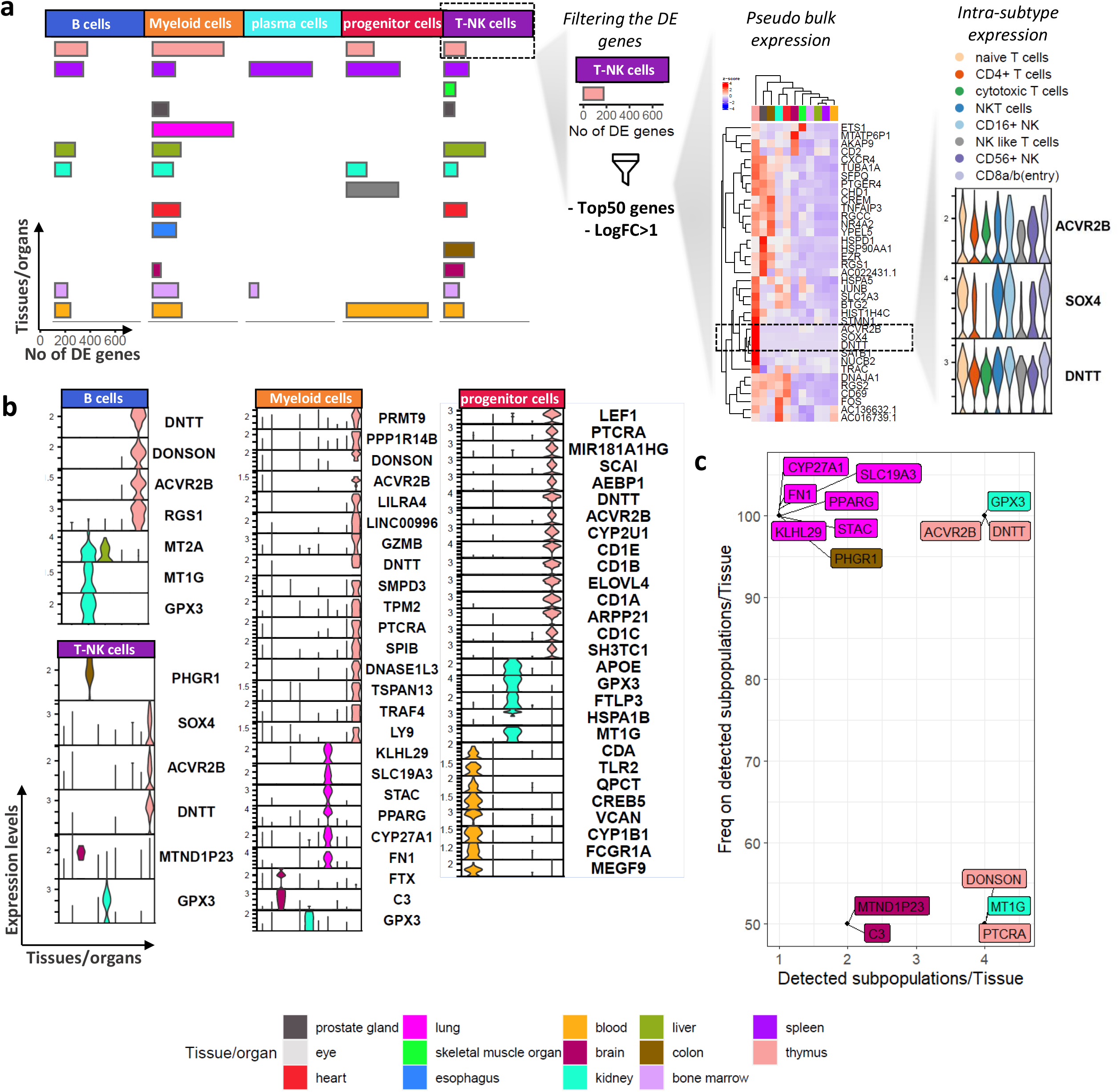
Organ-related signatures per immune subpopulations. **a,** Graph explaining the definition of the gene signature per organ in each immune cell type. The barplot shows the number of differentially expressed genes (overexpressed genes) in each organ splitting by the defined immune cell types. A filtering for the significantly overexpressed genes was applied and a hierarchical clustering analysis was performed using the top 50 genes. Finally, an intra-immune subtype expression analysis was performed checking the specificity of the expression. **b**, Violin plots showing the expression of the gene signatures per immune subtype and in each organ. **c**, Scatterplot showing the frequency of the detected immune cell subpopulations containing the gene in their signatures (x-axis) and the number of the detected immune cell types in each organ (y-axis).

Focusing on B cells, seven signature genes were uniquely overexpressed in specific organs (*DNTT, DONSON, ACVR2B* and *RGS1* in thymus, *MT2A* in liver, and *MT1G* and *GPX3* in kidney) (**Fig. 3b, Supplementary-Fig. 6b**). Interestingly, *RGS1* and *ACVR2B* genes in the thymus, are related to cytokine signaling further supporting their significance in the homing of different organs^12^, ^13^. Additionally, *GPX3* gene was uniquely expressed in kidney in line with previous publications mentioning its high expression in proximal tubules^14^.

The T-NK cell analysis revealed six genes (*PHGR1, SOX4, ACVR2B, DNTT, MTND1P23, GPX3*) uniquely expressed in specific organs (**Fig. 3b, Supplementary-Fig. 6c**). *PHGR1* was expressed in the colon, *SOX4, ACVR2B* and *DNTT* in thymus, *MTND1P23* in brain and *GPX3* in kidney. *PHGR1* associated with the colon is characterized by unique expression in the gastrointestinal tract^15^. Moreover, *SOX4* and *DNTT* in thymus are related to T cell development since CD8a/b (entry) was present only in thymus. Focusing on the myeloid cells, 16 signature genes were found for thymus (*PRMT9, PPP1R14B, DONSON, ACVR2B, LILRA4, LINC00996, GZMB, DNTT, SMPD3, TPM2, PTCRA, SPIB, DNASE1L3, TSPAN13, TRAF4, LY9*), however, most of them were associated with the biology of the plasmacytoid DCs, which were the enriched subpopulation of the thymic myeloid cells. Also, we found six signature genes for the lung (*KLHL29, SLC19A3, STAC, PPARG, CYP27A1, FN1*) associated with the biology of lung macrophages, and two signature genes for the brain (*FTX, C3*) and one for kidney (*GPX3*) (**Fig. 3b, Supplementary-Fig. 6d**). Also, most of the cells derived from the lung expressed *PPARG* and *CYP276A1*, which are expressed in the lung-specific macrophages and alveolar macrophages.

In terms of the progenitor cells (lymphoid/ myeloid lineages), blood showed eight uniquely expressed genes (*CDA, TLR2, QPCT, CREB5, VCAN, CYP1B1, FCGR1A, MEGF9*) followed by 15 genes in thymus (*PTCRA, MIR181A1HG, SCAI, AEBP1, DNTT, ACVR2B, TCF7, CYP2U1, CD1E, CD1B, ELOVL4, CD1A, ARPP21, CD1C, SH3TC1*) and five genes (*APOE, GPX3, FTLP3, HSPA1B, MT1G*) in the kidney (**Fig. 3b, Supplementary-Fig. 6e**). Since the thymus is enriched in a specific maturation process, its signature genes were associated with T cell differentiation, such as *CD1E* (T cell marker). The signatures regarding the different organs and immune cell types as well as the etiology of the expected or not expected gene expression are presented in **Table 2**.

**Table 2.**
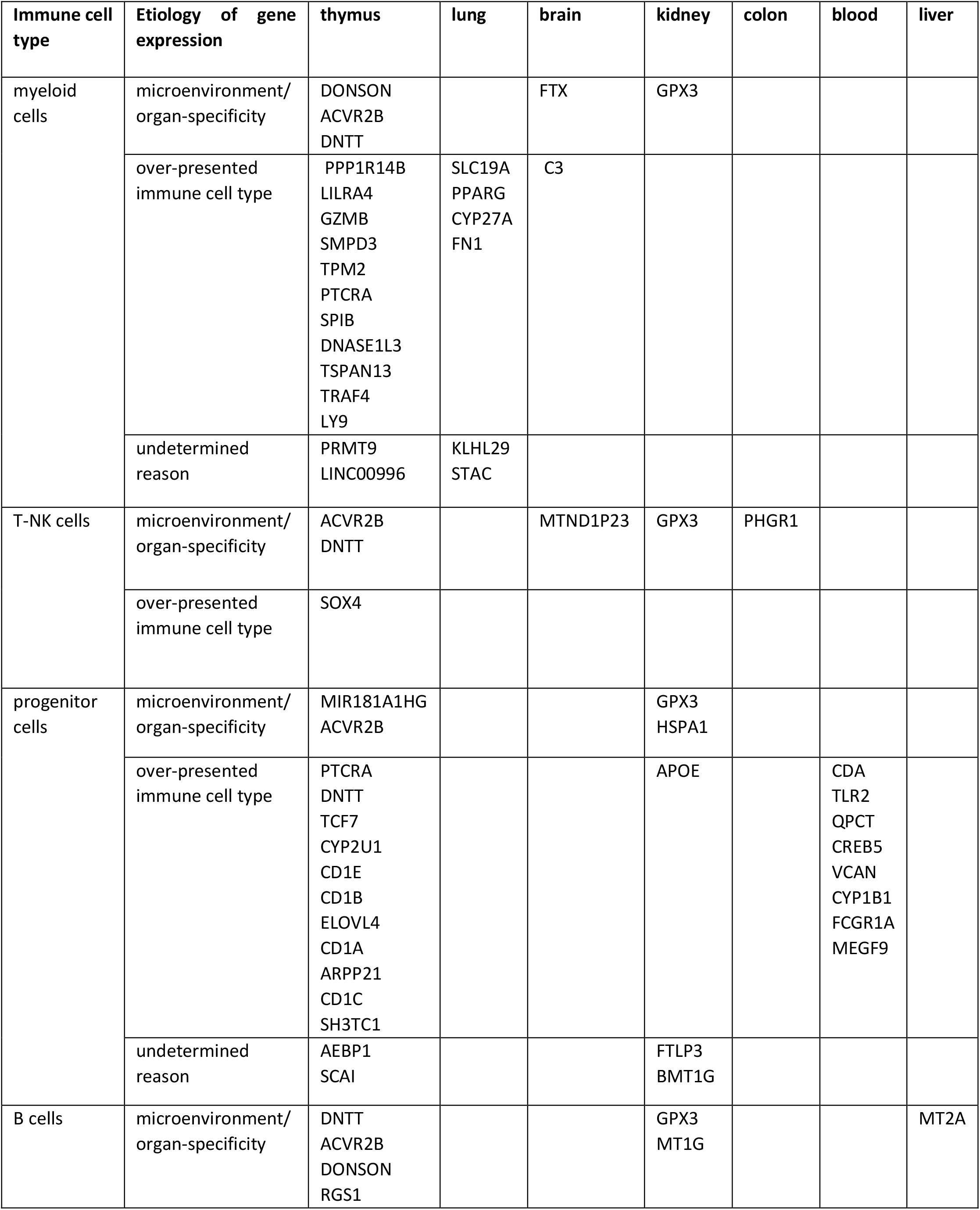
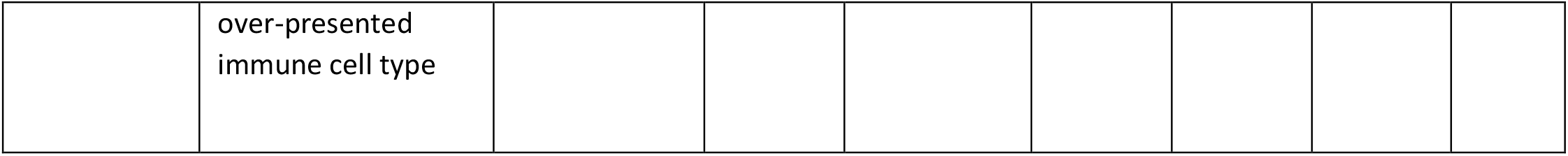
The organs’ gene signatures in each immune cell type adding the etiology of the detected expression as microenvironmental effect/ organ-specificity or influenced by the over-presented immune cell type. We defined as microenvironmental effects the expression of genes that are typically expressed in a particular organ and are not related to the immune cell type and as over-presented immune cell type etiology the genes that are typically expressed in a particular immune cell type. The genes that are not related to the previous categories were classified as undetermined reasons.

Signature genes are found in different immune compartments highlighting organ-specificity or the predominant immune cell type of each organ. They were not associated with intra-cluster immune cell subpopulations (**Supplementary-Fig. 6b-e**) and were expressed in a high proportion of cells (**Supplementary-Fig. 7**). Of all the signature genes identified in each immune cell subpopulation, some were unique to a specific immune cell subpopulation and organ, while others were shared across different immune cell subpopulations but exclusively expressed in one organ. Ten genes were present in more than 50% of the immune cell subtypes (B, T-NK, myeloid, progenitors) across all studied organs (**Fig. 3c, Table 2**). *GPX3* gene was a signature gene for kidney and *DNTT* and *ACVR2B* for thymus in all the immune cell types that were found on myeloid, progenitor, B and T-NK cells. The colon and lung showed signature genes that were only enriched in these organs, but only found in a single immune subtype since colon was enriched only in T-NK cells and lung in myeloid cells. These genes include *CYP27A1, FN1, KLHL29, PPARG, STAC, SLC19A3* for the lung and *PHGR1* for the colon. The revealed genes suggest a microenvironmental effect (*e*.*g. DNTT* in thymus, *GXP3* in kidney) or an expected observation based on the over-presented immune cell type in one organ (*e*.*g. PPARG* in macrophages derived from lung since it is a well-known marker for alveolar macrophages) (**Table 2**).

To validate our proposed signature genes for the thymus and kidney, we searched for additionally publicly available datasets. Focusing on the kidney, we found expression of *GPX3* in three independent scRNAseq datasets^16^, ^17^, ^18^(**Supplementary-Fig. 8a-b**). Regarding the thymic signature, the expression levels of *DNTT* and *ACV2RB* were validated using bulk RNA-seq from sorted *CD19*+ thymic cells^19^ (**Supplementary-Fig. 8a, c**).

### *In silico* functional analysis revealed transcription factor specificity and *NFkB* implication on localization

Going one step of organization further, we investigated the transcription factor (TF) activity (regulons) underlying expression pattern differences in different organs and immune compartments. The analysis was focused on the regulons that were uniquely found in one organ and were expressed in different immune cell subpopulations underlying the organ-specificity (**Fig. 4a**). Interestingly, eight regulons showed organ-specificity and were found in all the defined subpopulations *e*.*g. HNF4A* for the kidney and *TFF3* for the colon which have been previously described for these organs^20^, ^21^. Focusing on the regulons that appeared in more than three different organs, we found *PARG, POU2AF1, HOXB2* and *ESR2* (**Supplementary-Fig. 9a**). We compared the organ-specific signature genes (**Fig. 3b**) with organ-specific regulon target genes, resulting in four regulons associated with nine signature genes (**Table 3**). Of these four *HNF4* and *ZEB1* regulate more than one gene (**Fig. 4b**). *HNF4* was associated with *GPX3* and *MT1G* in B and progenitor cells from kidney and *ZEB1* with *TCF4, CD1A, ARP21* and *SH3TC1* in thymic progenitor cells.

**Fig. 4.**
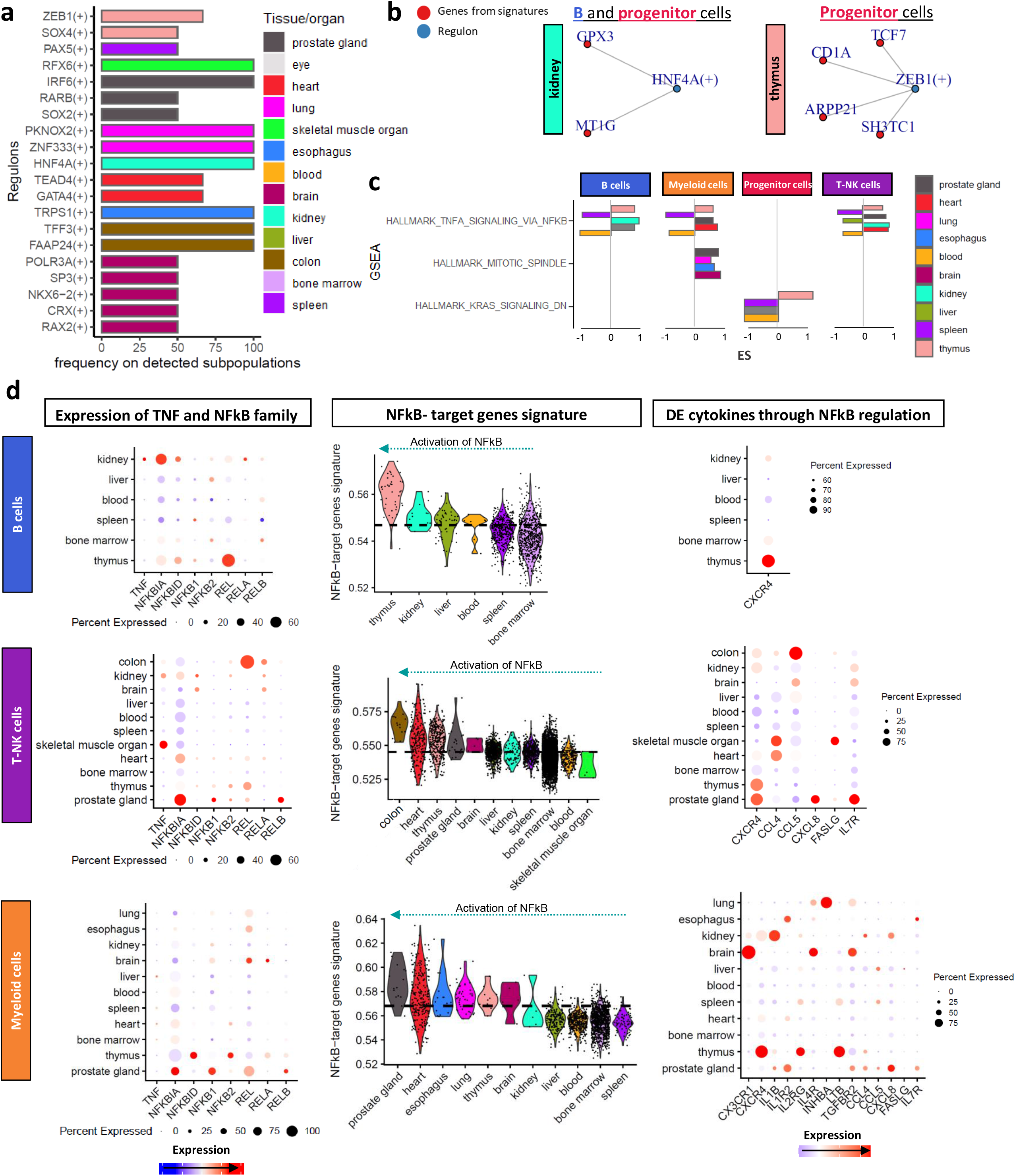
Functional analysis of differentially expressed genes across the different organs in each immune cell type. **a,** Barplot showing the unique regulons in each organ (y axis) and the percentage of the detected immune cell subpopulations showing the regulons in each organ (x axis). **b**, Network of the regulons (blue) and the genes associated with the gene-signature (red). **c**, Barplot showing the shared pathways in the GSEA (y axis) per immune cell type and in each organ and the enrichment score values (x axis). **d**, Analysis of the TNFA signaling pathway via NFkB in each immune cell type showing the expression of the TNF and NFkB family members, the target genes and the target cytokines.

**Table 3.** Organ-specific TFs in relation to the genes of the defined signatures.

| Regulons | Cell type | Organ | Gene in signature |
| --- | --- | --- | --- |
| CRX | myeloid_cells | brain | C3 |
| HNF4A | myeloid_cells | kidney | GPX3 |
| TFF3 | T-NK_cells | colon | PHGR1 |
| HNF4A | T-NK_cells | kidney | GPX3 |
| ZEB1 | progenitor_cells | thymus | TCF7 |
| ZEB1 | progenitor_cells | thymus | CD1A |
| ZEB1 | progenitor_cells | thymus | ARPP21 |
| ZEB1 | progenitor_cells | thymus | SH3TC1 |
| HNF4A | progenitor_cells | kidney | GPX3 |
| HNF4A | progenitor_cells | kidney | MT1G |
| RELA | B_cells | liver | MT2A |
| HNF4A | B_cells | kidney | MT1G |
| HNF4A | B_cells | kidney | GPX3 |

Using DE genes across immune subtypes and organs, GSEA (FDR<0.01) identified 12/26 uniquely enriched pathways in different organs (**Supplementary-Fig. 9b**). Established pathways such as WNT signaling pathway in thymic progenitor cells^22^ and ‘allograft rejection’ in blood B cells^23^ validates the analysis. Metabolic pathways were deregulated in immune compartments, particularly in the myeloid, with hypoxia being the most frequently affected across organs (spleen, heart, and kidney) (**Supplementary-Fig. 9b**). *TNF-α* signaling via *NFkB*, mitotic spindle, and *KRAS* signaling were targeted in more than three organs (**Fig. 4c**).

*TNF-α* signaling via *NFkB* was found in T-NK, myeloid, and B cells and is involved in normal cellular functions such as inflammation and immune cell interactions^24^, ^25^. We conducted an analysis by aggregating the genes associated with the *TNF-α* signaling via *NFkB* pathway from each DE analysis of each immune cell type, revealing a distinct gene set in each immune cell type across different organs (**Supplementary-Fig. 9c**). *TNFA* and *NFkB* family member expression varied across organs and immune cell types (**Fig. 4d**). *REL* was highly expressed in B and T-NK cells in the thymus, while *NFKBID* and *NFKB2* were expressed in the myeloid compartment. Uncommon immune cell types in specific organs showed high activation of *NFkB* target genes. For instance, B cells from the thymus showed higher activation compared to B cells from spleen or bone marrow (**Fig. 4d**). Additionally, the activation levels of the *NFkB* target genes were independent of the intra-subtype expression (**Supplementary-Fig. 9d**). Notably, the *NFkB* target gene activation in the maturation stage^26^ is expected, but we observed additional activation of the memory and mature memory cells from thymus and kidney compared to spleen which in turn showed enrichment in mature cells. We found 50 DE target genes for B cells, 109 for T-NK cells, and 274 for myeloid cells by performing differential expression analysis of the detected target genes across organs and immune cell types (**Supplementary-Fig. 9e**).

To further investigate the role of *NFkB* in cytokine activation and immune cell positioning^27^, ^28^, we checked for DE target cytokines across different organs and immune compartments (**Fig. 4d**). A different set of cytokines was found in T-NK cells (*CCL4, CCL5, CXCL8, CXCR4, FASLG, IL7R*) and myeloid cells (*CX3CR1, CXCR4, IL1B, IL1R2, IL2RG, IL4R, INHBA, LTB, TGFBR2*) combined with different expression profiles across organs. *CXCR4* was the only cytokine that overlapped between different immune compartment and was associated with thymic immune cells. Also, different cytokines were related to different organs *e*.*g. IL7R* in T-NK cells from the prostate gland and *CX3CR1* in T-NK cells from the brain. We examined cell-cell communication in thymic B cells and cardiac T-NK cells, which showed strong *NFKB* pathway activity, to confirm the importance of cytokines in host communication. *CXCR4-MIF* and *CXCR4-TGFB1* was found shared between thymic B cells and T cells, while interactions between cardiac T-NK cells and myeloid cells via *CCL5-CCR1, CCL5-CCRL2*, and *CLL4-CCR1* was observed (**Supplementary-Fig. 9f-h**).

## Discussion

Recent studies have provided organ-atlases combined with in-depth characterization of cells but lacked biological relevance. We present an immune meta-atlas using multiple organs from 12 publicly available projects in healthy donors (1/12 healthy donors from a liver transplantation), enables efficient analysis of public datasets in line with open science and data reusability. Compared to previous studies^5^, ^6^, ^7^, ^8^, our analysis extends knowledge beyond the annotation of rare subpopulations and focuses on immune cell residency across different organs. We aimed to investigate the changes that immune cells undergo while residing in host organs, their use of components of the host organ, and their potential roles beyond their defensive function. Following previous publications^5^, ^29^, unsupervised analysis of multi-organs immune cells separated the distinct immune subpopulations showing no major differences in transcriptomic landscape across organs. We report certain immune cell clusters that are present across different organs, but also found organ-specific macrophages in line with previous publications^7^. The transcriptomic signatures of Tregs found in various organs suggest that these cells have undergone local adaptation to maintain immune homeostasis. Interestingly, Tregs in the spleen appear to have a thymic origin, while Tregs in other organs suggest differentiation in the periphery, rather than being generated in the thymus^30^.

Primary/secondary lymphatic organs showed the expected immune cell types distribution while non-lymphoid organs showed myeloid cell enrichment. Moreover, lung and esophagus had non-correlated patterns due to specific macrophage enrichment. We observed a novel correlation between immune cells from the thymus and kidney, with good correlation across different immune compartments. Interestingly, we found that progenitor cells from the kidney were clustered with classical thymocytes, indicating a thymocyte-like phenotype. This relationship is intriguing given the previous association between thymus and kidney diseases, though the underlying mechanism remains unknown^31^, ^32^, ^33^. Additionally, we found evidence to suggest that thymic B cells originate from the bone marrow, as their transcriptome showed the strongest correlation with bone marrow cells compared to blood and spleen cells^34^. Even though the immune cells are consistent across different organs, the upregulation and downregulation of genes observed in the study could be attributed to organ-specific gene expression programs. This highlights the requirement for immune cells to adapt to the specific microenvironment of each organ and carry out specialized functions.

Using hierarchical clustering method based on the DE genes (upregulated and downregulated genes), we detected only certain upregulated genes showing unique expression per organ and immune compartment and these gene signatures were independent of the intra-subtype expression. Some of these upregulated genes were present in all the detected immune cells type per organ suggesting the usage of specific factors or molecules present in the host organ microenvironment (*e*.*g. DNTT* and activin receptor, *ACVR2B*, in thymus, *GXP3* in kidney). Other genes with high organ-specificity were expressed only in one cell type without association with the host organ highlighting an expected observation based on the over-presented cell type in one organ (*e*.*g SOX4* expression in thymus T cells came from thymocytes differentiation^35^, *PPARG* in macrophages derived from lung which is a marker for alveolar macrophages^36^). Focusing on the microenvironmental effect of host organs, the immature stage of B and T cells express *DNTT*, but the maturation events require its silencing^37^, ^38^. Here, we report a high percentage of cells located in thymus also expressed *DNTT*, which may be associated with microenvironmental signals. Kidney-derived immune cells expressed GPX3, an exclusive kidney antioxidant enzyme that protects high metabolic activity cells from oxidative stress, as shown in previous studies on macrophages during kidney disease^14^, ^39^. Regulons analysis associated *GPX3* with *HNF4A* which is a TF associated with the kidney pathophysiology^20^.

We found deregulation of metabolic pathways which supports that immune cells altered them to be able to enter and adapt to the environment of the host organ contributing to homeostatic tissue function^40^. The most targeted pathway was *TNFa* signaling via *NFkB* showing different *NFkB* activity through distinct expression patterns of *NFkB* family members and different expression profiles of the target genes in various organs. Additionally, we observed an activation of the NFkB target genes in specific organs where such immune cells are not commonly found (*e*.*g*. B cells in thymus) analyzing the data in each immune compartment and may provide evidence of a critical role of *NFkB* in homing. Previous publications showed the implication of *NFkB* in localization since regulates migratory dendritic cells^41^ and new T cells and recent thymic emigrants up-regulated IL-7Rα once they leave the thymus which is a target gene of the *NFkB* transcription network^42^. Knowing its role in the transcriptional activation of cytokines, we found different repertoires of cytokines through *NFkB* regulation in different organs further supporting the role of *NFkB* in cell positioning through cytokines^27^.

Taken together, this study reports organ-specific genes in immune cells due to the microenvironment of the host organs. Unique and shared regulons and pathways were identified underscoring the functional impact of the altered expression profiles of the immune cells through different organs. The analysis demonstrated the implication of the *NFkB* pathway in cell positioning. Alterations in gene expression and metabolic pathways implying immune cells are not only involved in defending host organs but also adopt the host environment and contribute to their homeostasis and repair. The ability of immune cells to sense and respond to changes in the environment of different organs further highlights the importance of understanding the interplay between immune cells and host organs in health and disease.

## Methods

### Data collection and aggregation

162 loom files were downloaded from the 12 different project of the HCA consortium (**Supplementary Table 1, Supplementary methods**). Of note, one project was using healthy donors from liver transplantation including samples from liver, blood and spleen (**Supplementary Table 1**). The files were transformed into Seurat objects with the function as.Seurat() from Seurat^43^ package in R. After incorporating metadata information, each object was screened for immune cells based on the expression of the *CD45* immune marker. Specifically, cells were considered immune cells if they exhibited at least one read of the *PTPRC* - *CD45* surface marker gene. This selection process resulted in a reduction of immune cells within each file, particularly in the case of bone marrow, where up to 80% of cells were excluded, primarily from the progenitor lineage. The individual files were merged into a single one. The full script of the process is available at GitHub (https://github.com/biomedicalGenomicsCNAG/mulTI_Metalas).

### Quality control, batch effect correction and clustering

We performed all downstream pre-processing with Seurat^43^. For quality control, we excluded cell barcodes with <1,000 Unique Molecular Identifiers (UMI), <200 detected genes, or mitochondrial expression >15%. In addition, we excluded genes detected in <=4 cells. Next, we applied the functions NormalizeData(), FindVariableFeatures(), ScaleData() and RunPCA() of Seurat (with default parameters). Batch effect correction was performed by applying Harmony^44^ with the top 30 Principal Components (PC) as input. We considered as batches the cells coming from (i) the same sample and (ii) the same subproject of the HCA consortium. The validation of the data integration was performed visually with UMAP using the function RunUMAP() on the first 30 PCs and quantitatively with the Local Inverse Simpson’s Index using the LISI package in R (**Supplementary-Fig. 1a**).

To cluster cells into groups, we used the function FindNeighbors(reduction = “harmony”, 30 PCs) and then determined the clusters based on the function FindClusters(). Low-quality clusters (*e*.*g*., high expression of mitochondrial or rRNA, doublets) were removed from the downstream analysis. The characterization of the cells was performed by subsetting and reapplying FindNeighbors(reduction = “harmony”, 30 PCs) and FindClusters().

### Downstream analysis

Cell annotation was performed manually using the most significant markers in each cluster as well as comparing the results with *SingleR* and *EnrichR* tools in R. A publicly available annotation of the tonsil was used for in-depth characterization of the immune applying *Azimuth*^9^. Differential expression analysis was performed considering as a threshold the presence of at least 40 cells in each analyzed group. Except for the default parameters of FindMarkers() we additionally used as a criterion the minimum percentage of cells expressing one gene to be greater than 50% on the group of interest helping to avoid the detection of genes that expressed only in a small proportion of cells. A pseudobulk table was generated using the function AverageExpression() from the Seurat package. UCell package^45^ was used to calculate the expression gene signature of *NFkB* target genes per cell using the AddModuleScore_UCell() function. Gene set enrichment analysis (GSEA) for each gene list was performed using the database of MsigDB under the package *clusterProfiler*^46^ in R. To infer transcription factor (TF) activity (known as regulons), we used *pySCENIC* on the scRNA-seq raw matrices from the different organs and immune cell types. The regulon activity per cell was quantified using an enrichment score for the targets of each regulon (AUCell). Additionally, regulon specificity score (RSS) was computed using the Jensen-Shannon Divergence^47^. Pseudotime analysis was performed using *slingshot* and cell-cell communication using *LIANA* packages in R. All the visualization was performed using the *Seurat, ggplot2, ComplexHeatmap* and/or *corrplot* package in R.

### Data Sharing Statement and Code availability

All the loom files were downloaded through HCA portal (**Supplementary Table 1**). All the scripts for the analysis of the loom files until the visualizations are available on Github (https://github.com/biomedicalGenomicsCNAG/mulTI_Metalas) with a detailed description of each one. scRNA-seq expression matrix is available at Zenodo (https://doi.org/10.5281/zenodo.7756209).

## Acknowledgements

This research has received funding from the European Union’s Horizon 2020 research and innovation programme through the ERC Synergy project BCLL@lasunder grant agreement No 810287 (I.G.G.). M.T. was granted with the ‘Seal of excellence ISCIII-HEALTH’. Institutional support was from the Spanish Instituto de Salud Carlos III, Fondo de Investigaciones Sanitarias and cofunded with ERDF funds (PI19/01772). We acknowledge the institutional support of the Spanish Ministry of Science and Innovation through the Instituto de Salud Carlos III and the 2014–2020 Smart Growth Operating Program, to the EMBL partnership and institutional co-financing with the European Regional Development Fund (MINECO/FEDER, BIO2015-71792-P). We also acknowledge the support of the Centro de Excelencia Severo Ochoa, and the Generalitat de Catalunya through the Departament de Salut, Departament d’Empresa i Coneixement and the CERCA Programme to the institute. We thank Miranda Stobbe, Maria Terradas and Jose I. Martin-Subero for revising the manuscript. Also, we would like to thank the generator of the used dataset that were accessible through HCA portal.

