## Supplementary methods, Figures and Table for "Multi-organ single-cell transcriptomics of immune cells uncovered organ-specific gene expression and functions"

### Supplementary Methods
#### Study group
162 loom files were downloaded from the 12 different projects of the HCA consortium. In more detail, we focused on 14 different organs: prostate gland[1, 2], eye[3], heart[2, 4], skeletal muscle organ[2, 5], blood[6], liver[6], spleen[6], brain[7, 8], kidney[9], colon[10], esophagus[2], lung[2], thymus[11], bone marrow (1M Immune Cells project).


### Supplementary Figures
**Supplementary Figure 1. a,** Boxplot showing the iLISI scores for uncorrected and corrected values after running Harmony concerning the different tissues. **b,** Barplot showing the number of samples that were available for each organ. **c,** UMAP showing the expression levels of *CD45* through different

immune compartments highlighting the lack of the non-thymic progenitor cells. **d,** Barplot showing per immune cell type the distribution of the organs. **e,** The UMAP plot of the immune cells is colored by the organ from which they came. **f,** The UMAP plot of the immune cells colored by the cell cycling markers.

**Supplementary Figure 2.** Heatmap showing the -log10(p-value) of the hypergeometric distribution between the annotation of tonsil atlas[12] and the different organs. The asterisks indicate statistically significant enrichment p-value<0.05 (hypergeometric distribution).

**Supplementary Figure 3. a,** Expression levels and percentage of cells expressing the gene markers. **b,** Proportions of the organs in each B cell type. **c,** Correlation (Pearson's r) plot between the expression profiles of B cells in different organs. **d,** Correlation plots between the expression profiles of mature memory (left) and memory B cells (right) subpopulation in different organs which include more than 10 cells. **e,** Expression levels and percentage of progenitor cells expressing the gene markers. **f,** Proportions of the organs in each progenitor cell type.

**Supplementary Figure 4. a,** Expression levels and percentage of T-NK cells expressing the classical gene markers. **b,** Proportions of the organs in each T-NK cell type. **c,** The UMAP plot of the memory T cells. **d,** Heatmap showing the -log10(p-value) of hypergeometric distribution between memory T cell subpopulations and the different organs. The asterisks indicate statistically significant enrichment p-value<0.05 (hypergeometric distribution). **e,** The UMAP plot of the *CD4*+ memory B cells including Treg subpopulation. **f,** Expression levels and percentage of *CD4*+ and Treg cells expressing the gene markers. **g,** Barplot showing the number of Treg cells across the different organs. **h,** Heatmap of Tregs based on 356 DE genes across the different organs. **i,** Correlation plot of the expression profiles of T-NK cells in different organs.

**Supplementary Figure 5. a,** Expression levels and percentage of myeloid cells expressing the gene markers. **b,** Proportions of the organs in each myeloid cell type. **c,** Correlation plot of the expression profiles of myeloid cells in different organs. **d,** The UMAP plot of the macrophage subpopulations including the *CD14+* cells. The lines represent the trajectories using the slingshot package in R. **e,** The UMAP plot of the characterized macrophage subpopulations. **f,** Expression levels and percentage of macrophages expressing the gene markers. **g,** Proportions of the organs in each macrophage subpopulation. **h,** Violin plots showing the expression of the *CD79A* marker of B cells as well as the expression of the IGHG genes in different organs.

**Supplementary Figure 6. a,** Barplot showing the number of differentially expressed genes (downregulated genes) in each organ splitting by the defined immune cell types. **b-e,** Violin plots for each immune subtype showing the expression of the gene signatures in each organ and barplots showing the number of cells per immune subtype in **b,** B cells, **c,** T-NK cells, **d,** myeloid cells and **e,** progenitor cells.

**Supplementary Figure 7.** Violin plots for each immune subtype showing the expression of the gene signatures in each tissue and dotplots showing the expression levels and the percentage of cells in each tissue expressing the gene in **a,** B cells, **b,** T-NK cells, **c,** myeloid cells and **d,** progenitor cells.

**Supplementary Figure 8. Validation of gene signatures using additional datasets. a,** Expression levels and percentage of the immune cells expressing the *ACVR2B*, *DNTT*, *GPX3* genes in the present study. **b,** Expression levels and percentage of the immune cells expressing *GPX3* gene in three additional datasets serving as validation cohorts. **c,** The dotplot (left) shows the expression levels of *DNTT* and *ACV2RB* in bulk RNA-seq data from sorted *CD19*+ thymic cells. *BCL6* was used as a positive control gene and *GPX3* as a negative control. The boxplot (right) displays the expression levels of all genes after DESeq2 normalization. The red lines highlight the minimum level of expression of the DESeq2 normalization.

**Supplementary Figure 9. a,** Barplot showing the shared regulons across organs (y axis) and in which immune cell subpopulations they were detected (x axis), colored by organ. **b,** Barplot showing the pathways that are enriched according to the GSEA (y axis) per immune cell type and in various organs and the ES values (x axis). Arrows highlighting the metabolic pathways. **c,** Heatmaps showing the DE genes in each tissue in the *TNFA* signaling via *NFKB* pathway grouped by immune compartment. **d,** Violin plots showing the activation of NFkB targets genes in each immune cell type including a deeper annotation of the cells, **e,** Heatmaps showing the DE genes in each tissue regarding the target genes of NFkB grouped by immune compartment. **f-h,** Cell-cell communication analysis showing the interaction specificity and the expression magnitude for **(f)** B cells (Receptor) with T-NK cells (ligand), **(g)** *CXCR4*+ B cells (Receptor) with Tregs (ligand), **(h)** T-NK cells (ligand) with B cells (Receptor) and myeloid cells (Receptor).

#### Supplementary Figure 1

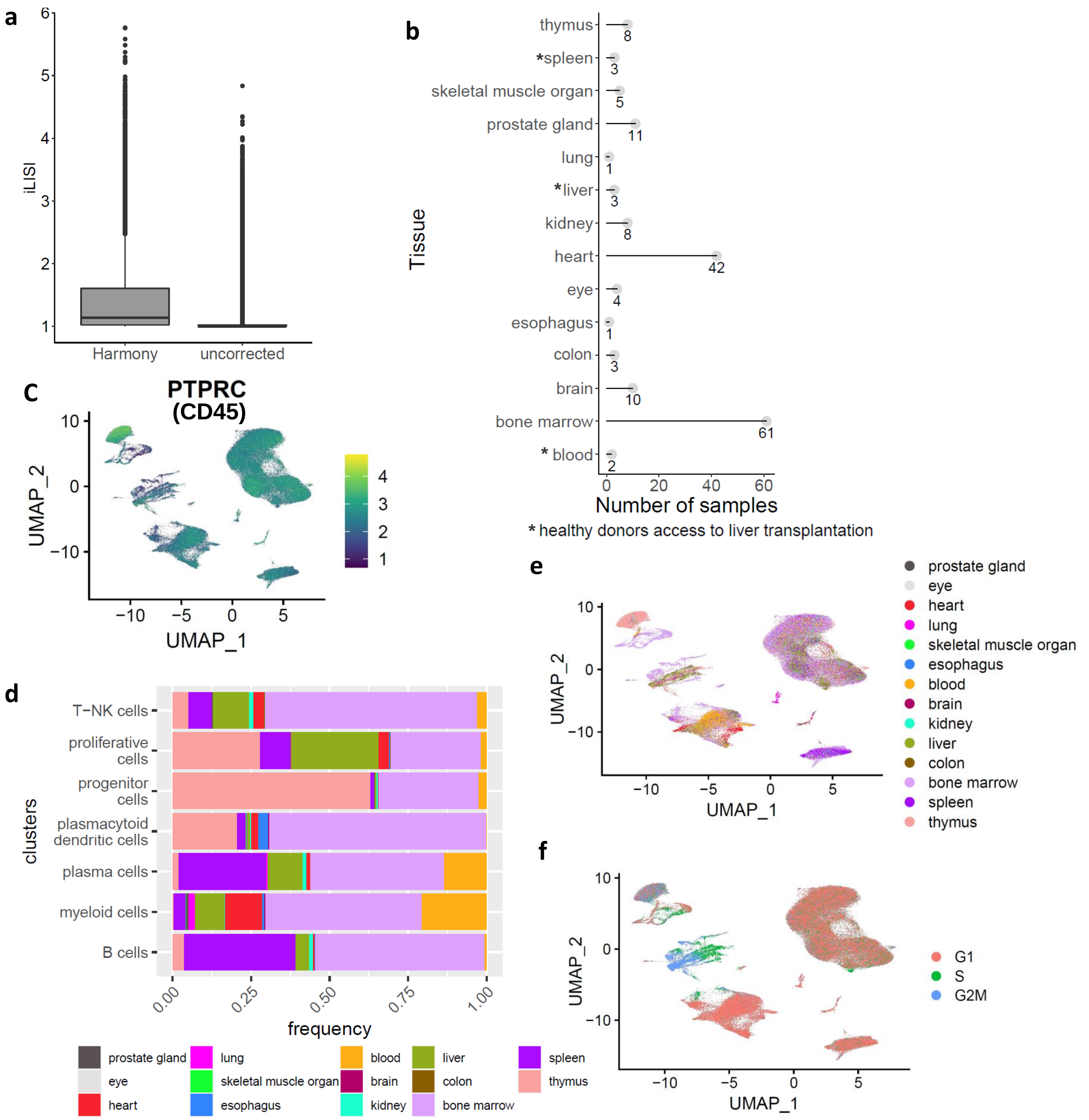

### Supplementary Figure 2

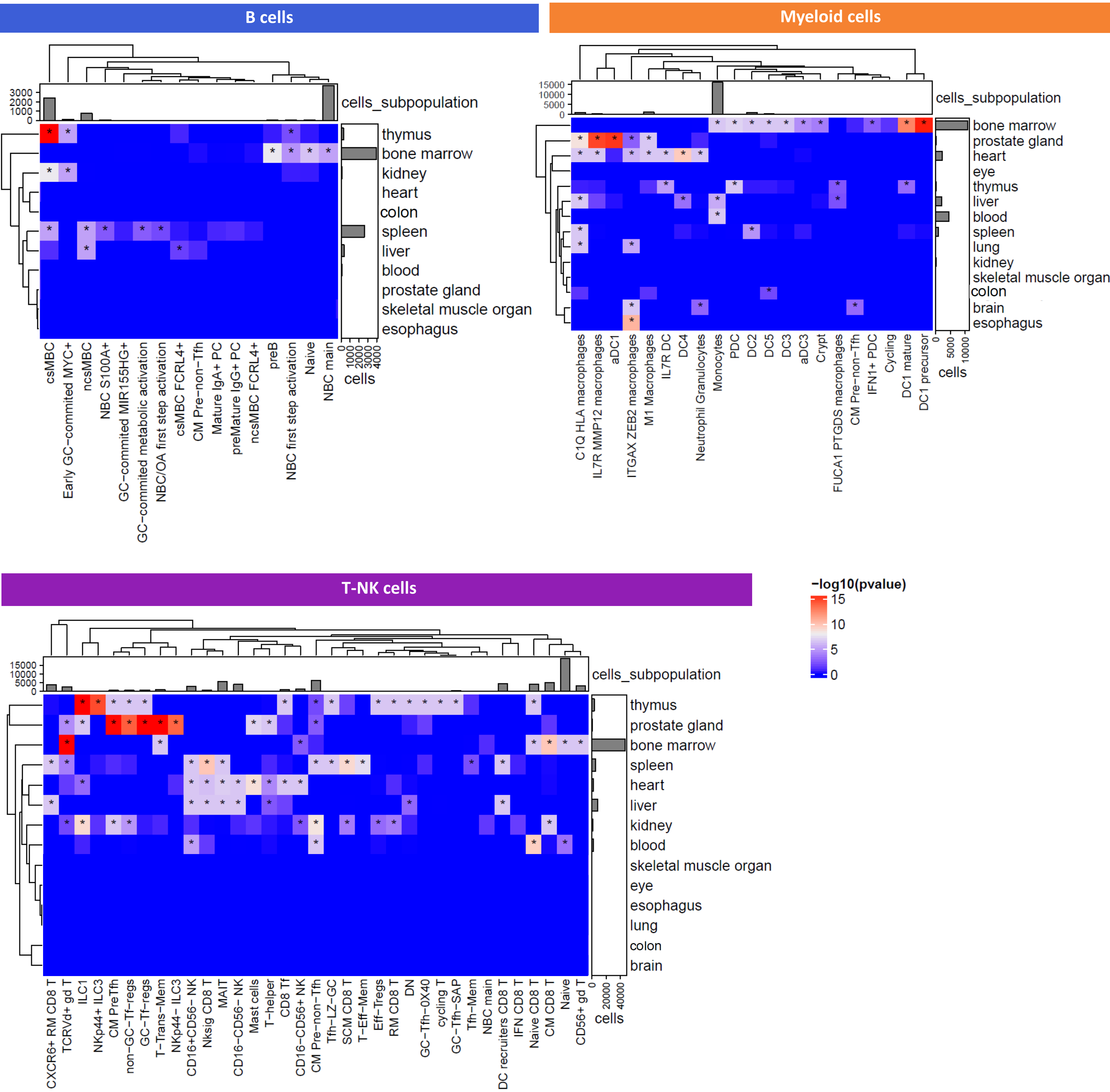

### Supplementary Figure 3

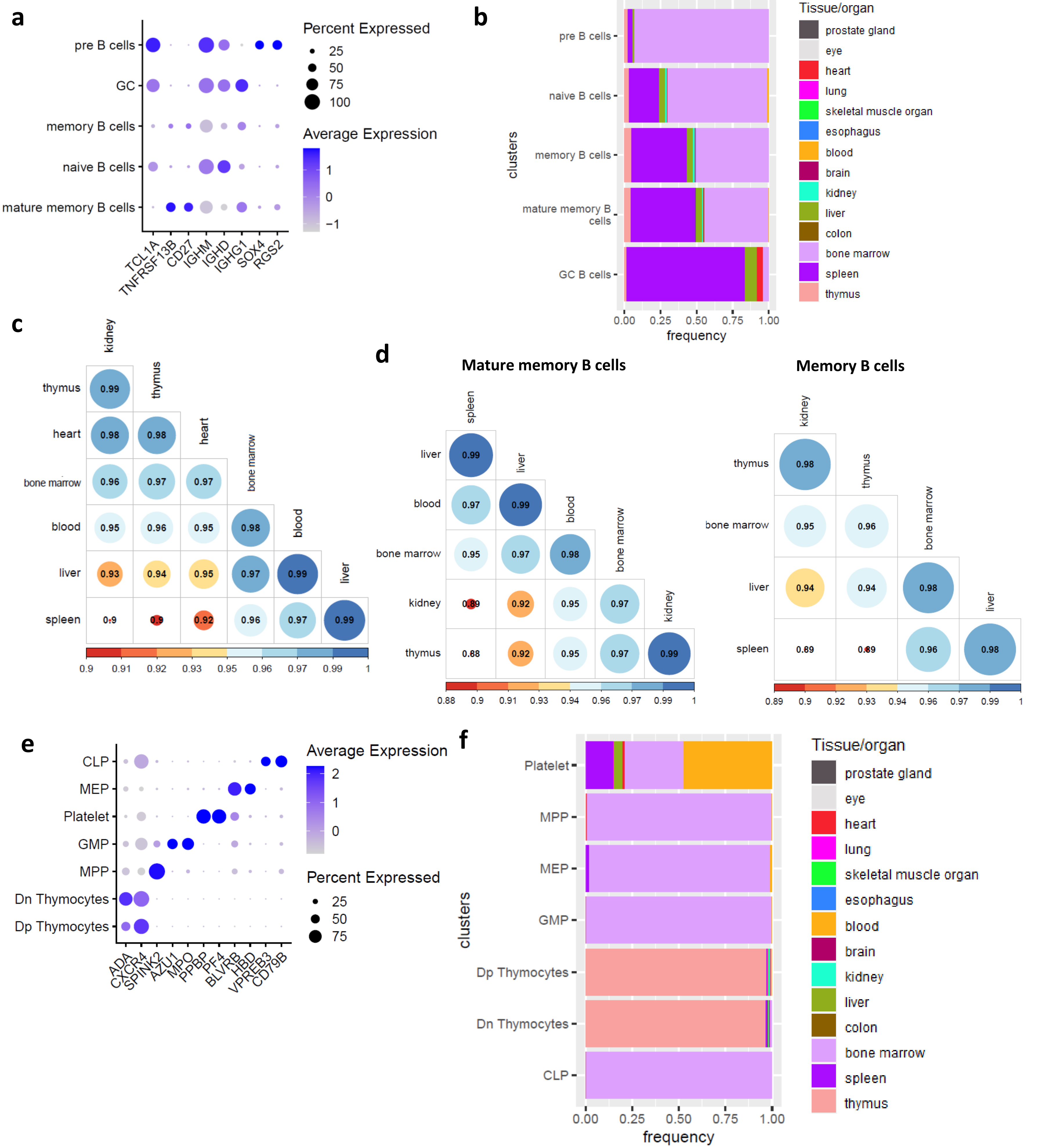

#### Supplementary Figure 4

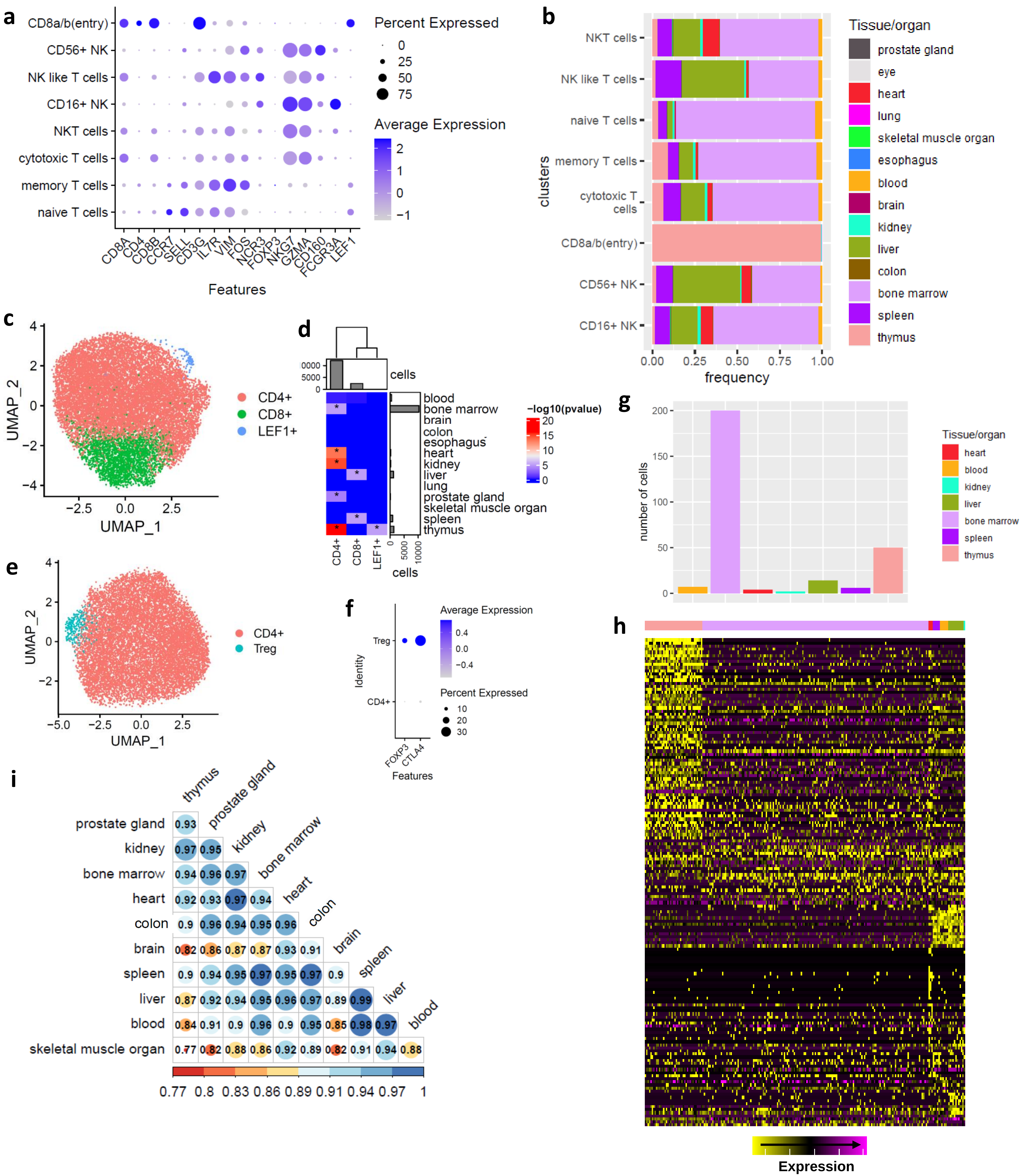

### Supplementary Figure 5

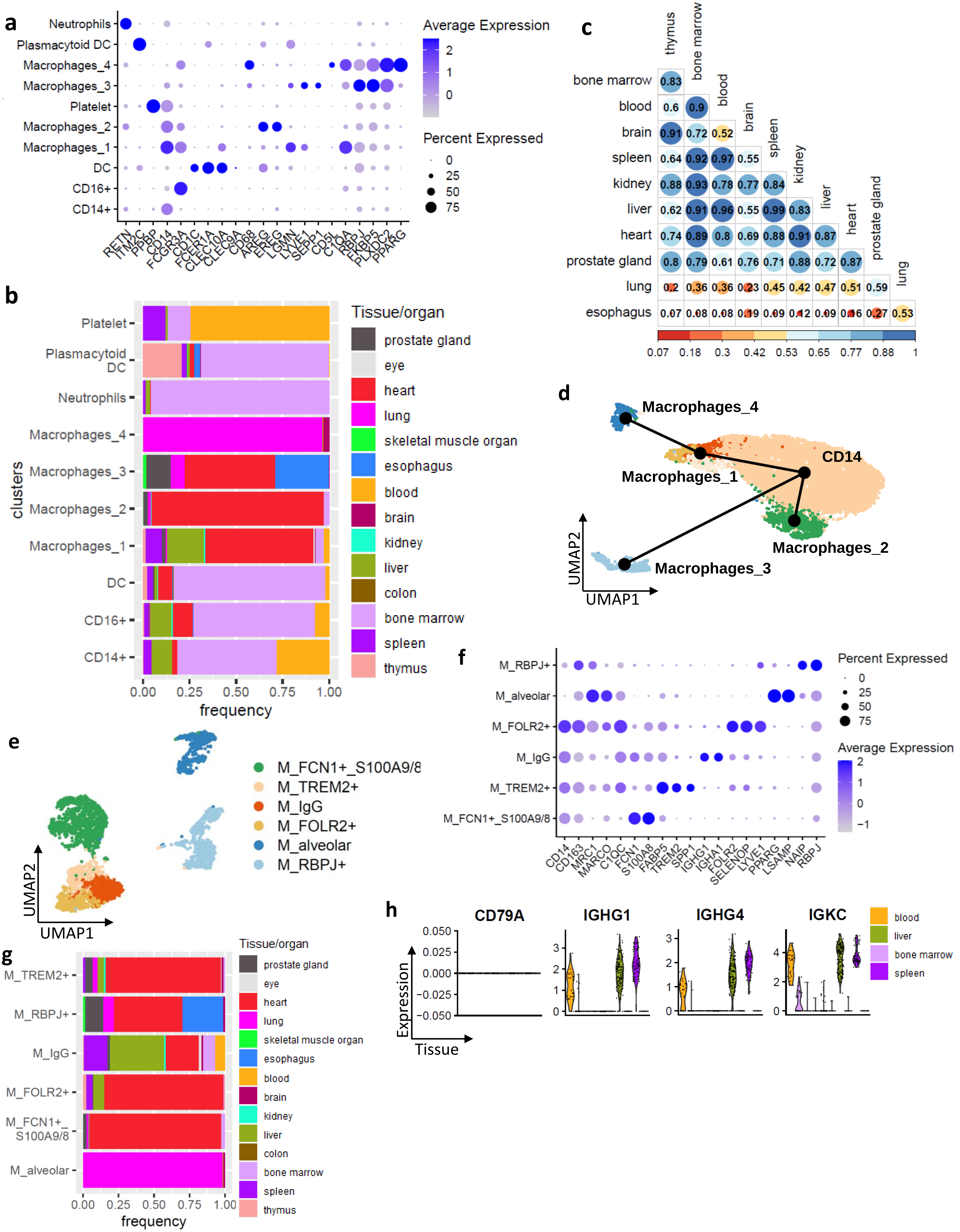

### Supplementary Figure 6

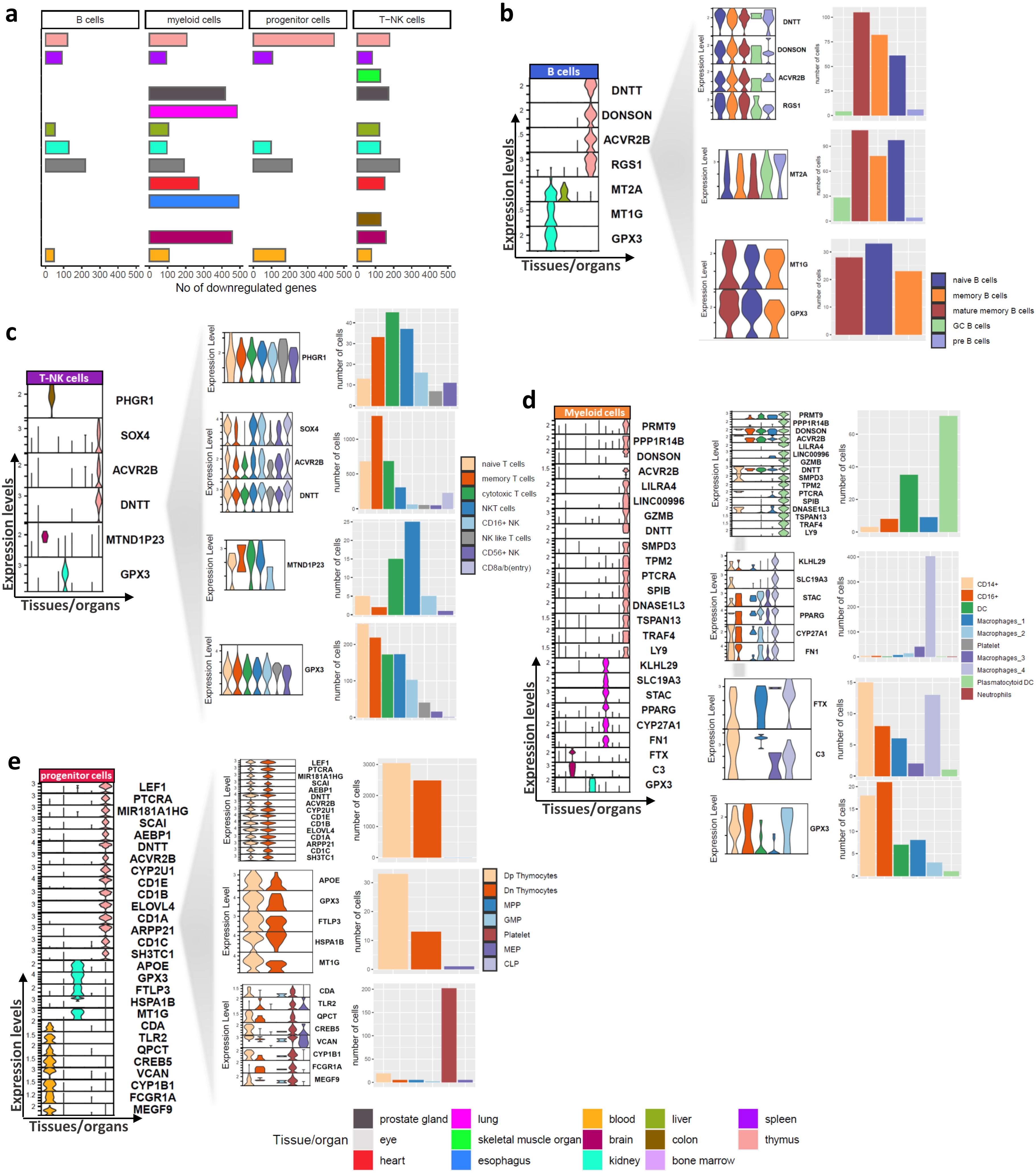

### Supplementary Figure 7

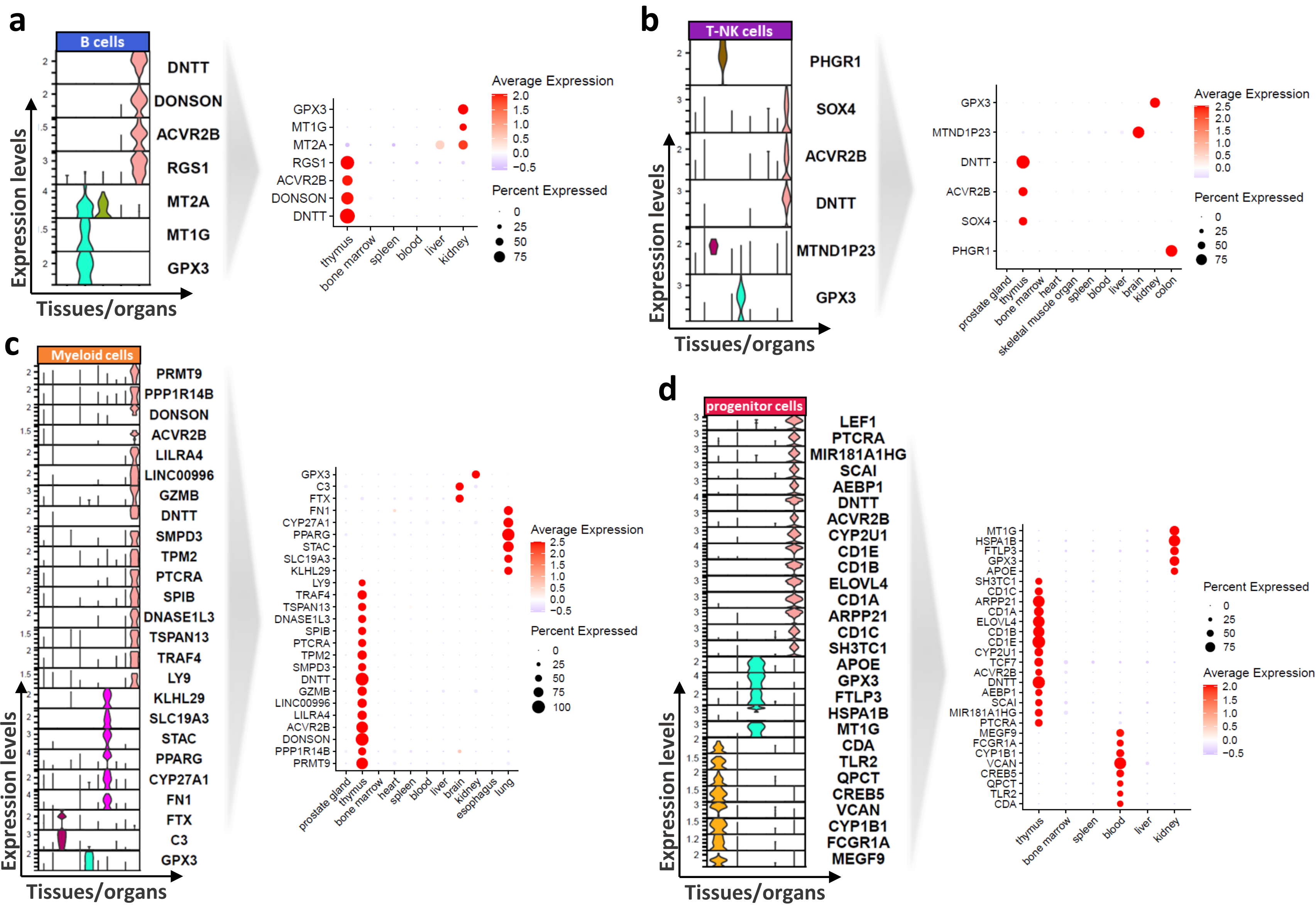

#### Supplementary Figure 8

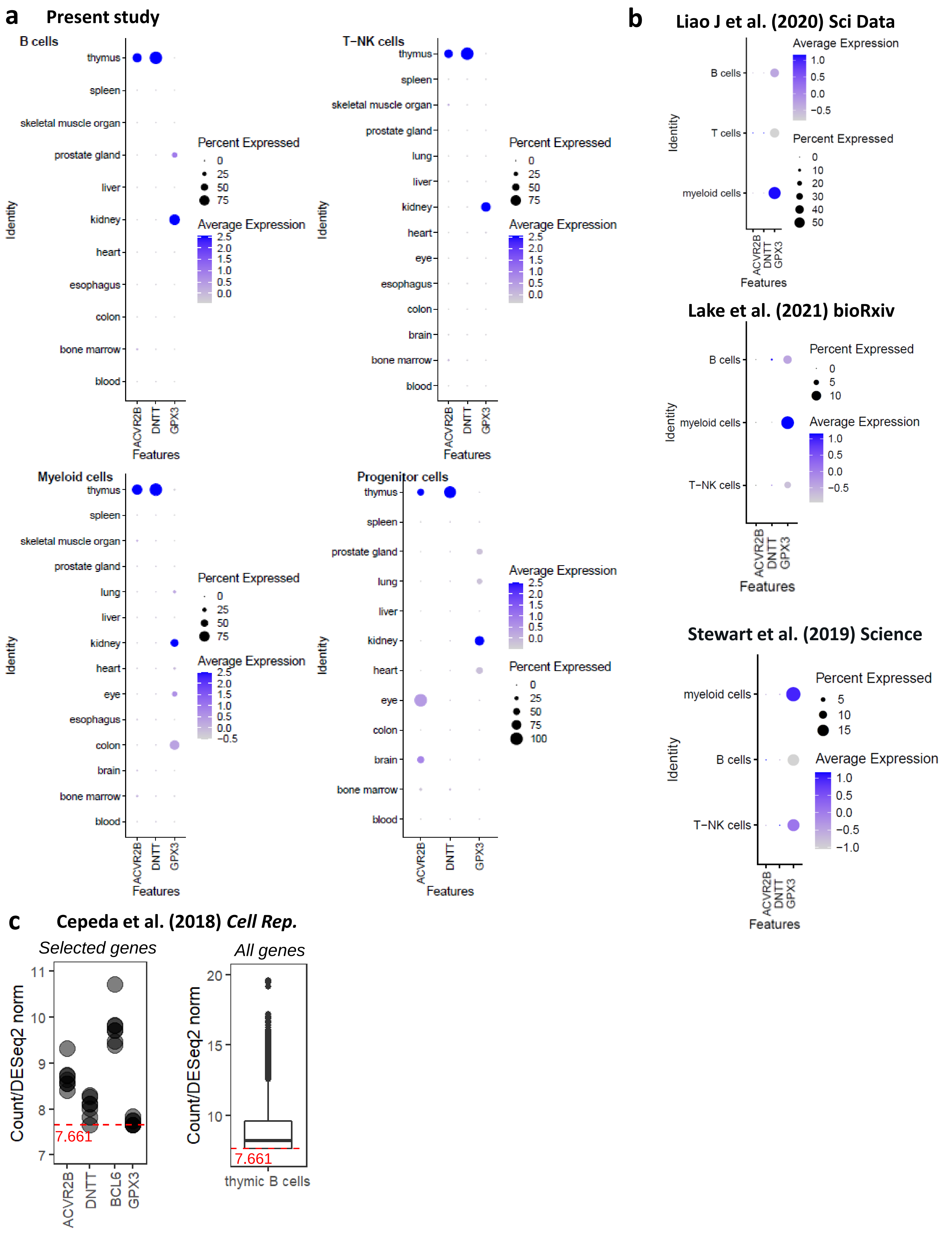

### Supplementary Figure 9

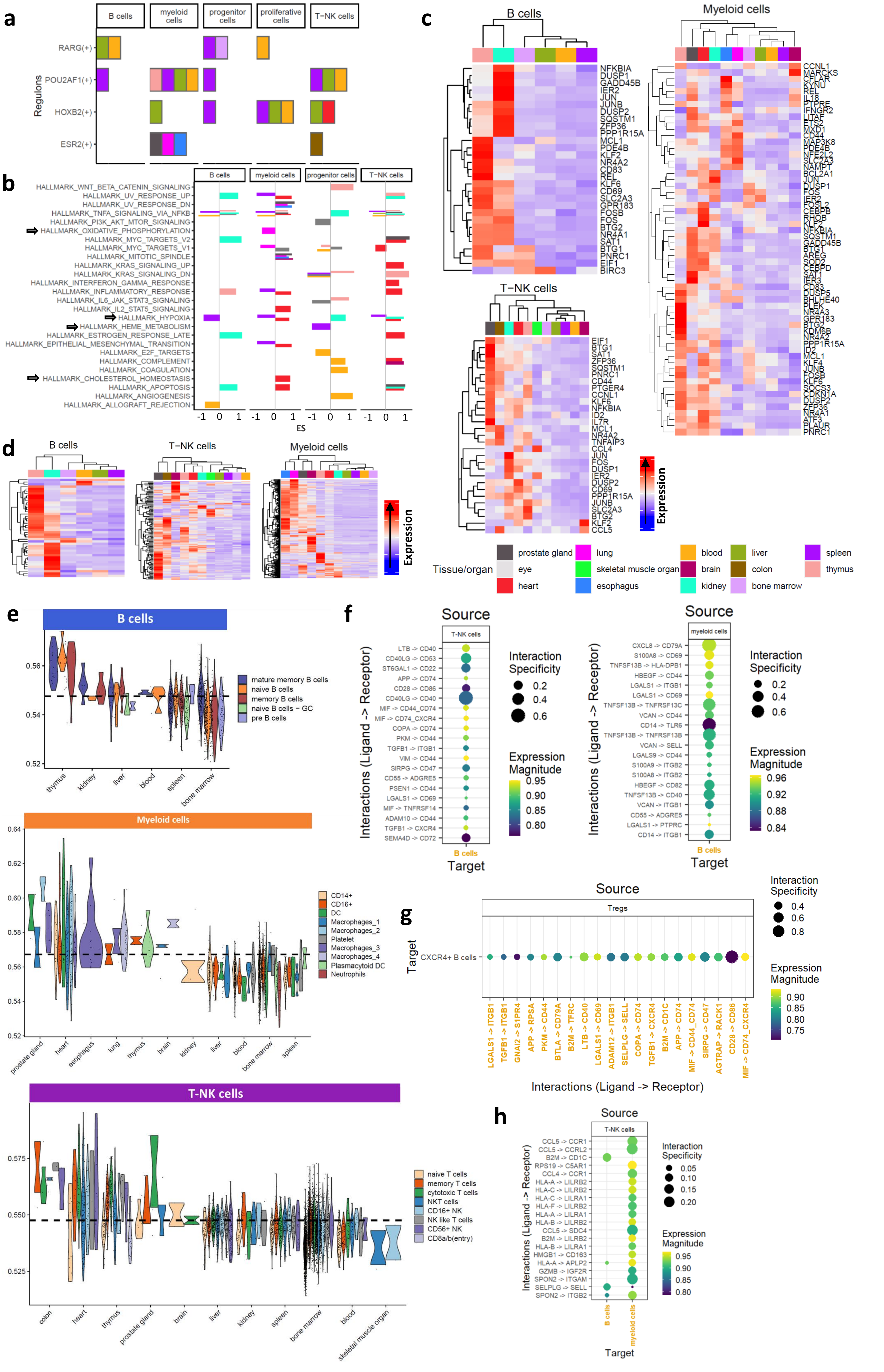

**Table S1. Study group**

| Sample_ID | Project | Tissue |
|---|---|---|
| 004c76c1-959c-5b0b-b8dc-f9812acc4778 | 1M Immune Cells | bone marrow |
| 09839dc3-7d1e-5f89-812f-bee5c2586a16 | 1M Immune Cells | bone marrow |
| 154e684c-9c89-5456-8424-7dad0d381967 | 1M Immune Cells | bone marrow |
| 168757c2-1cae-53a0-af63-d9a89b9aa417 | 1M Immune Cells | bone marrow |
| 1d630a02-bd52-5ac1-8ec6-6572de2d582a | 1M Immune Cells | bone marrow |
| 1dfb563b-2095-5290-b9cf-9c2c19ed3913 | 1M Immune Cells | bone marrow |
| 2662cfa0-fce5-5d80-8f24-f6c760507ecd | 1M Immune Cells | bone marrow |
| 2900c8a8-01c3-52f7-9375-1b0f98b15738 | 1M Immune Cells | bone marrow |
| 3ac6afe2-69ae-5535-9f5e-1529e9b6ca40 | 1M Immune Cells | bone marrow |
| 3ba67df1-837b-5515-a8a5-a834ba4560bb | 1M Immune Cells | bone marrow |
| 3f0ce691-a96f-56a2-b983-8bb048d052ea | 1M Immune Cells | bone marrow |
| 434867a6-1279-5906-b59e-1b8e0dfd99dc | 1M Immune Cells | bone marrow |
| 4e7f3a56-4888-521e-b8df-e5c06c4b24c1 | 1M Immune Cells | bone marrow |
| 568bac66-3ef8-52e1-a967-5c8c8100a924 | 1M Immune Cells | bone marrow |
| 58c4f886-924e-5877-a7fe-811ebfa79542 | 1M Immune Cells | bone marrow |
| 5bd755dd-4623-5b67-a0b0-1747bd6c6652 | 1M Immune Cells | bone marrow |
| 68448a63-5f1e-5926-9724-d63e875068a0 | 1M Immune Cells | bone marrow |
| 687e8127-583b-555f-a618-b603dbc757a8 | 1M Immune Cells | bone marrow |
| 7309180a-5924-542f-bf5a-e2c5234e5470 | 1M Immune Cells | bone marrow |
| 7adde500-de8d-51bb-8033-5a9b0dc503d0 | 1M Immune Cells | bone marrow |
| 83d50dd1-c7a8-52c2-ba43-456b2d4264e5 | 1M Immune Cells | bone marrow |
| 89bae042-beaf-5952-9212-23629385e886 | 1M Immune Cells | bone marrow |
| 9820765b-a8bb-5949-98c0-317c4353b185 | 1M Immune Cells | bone marrow |
| b5b28d72-5d7d-56f0-a253-ec514f3549eb | 1M Immune Cells | bone marrow |
| b876153e-b396-5904-9daf-0b445f24411a | 1M Immune Cells | bone marrow |
| c0bbea68-0956-5795-a6d6-30d494d42188 | 1M Immune Cells | bone marrow |
| c319ad92-85aa-53e7-b452-6d7eaa3a48b2 | 1M Immune Cells | bone marrow |
| c3c88d34-3b7b-55fd-83a8-fdb3f12f1af7 | 1M Immune Cells | bone marrow |
| c66731ea-0e30-57f8-b732-21e3de8b0cc7 | 1M Immune Cells | bone marrow |
| c737d21f-c3f0-52d9-ac07-c3329c99a7dc | 1M Immune Cells | bone marrow |
| c7affe2f-f1bb-5f45-af76-51589bd6587e | 1M Immune Cells | bone marrow |
| cf87010d-e2e9-5ad1-9ad5-7bcf2780f659 | 1M Immune Cells | bone marrow |
| e0ad2d93-9dfb-542f-8f93-ac2d2ed6f9f7 | 1M Immune Cells | bone marrow |
| f139fca6-7425-5440-b586-0240f76b6082 | 1M Immune Cells | bone marrow |
| 07d7019b-03f3-5ecf-a9fe-0a870e4c5f9b | 1M Immune Cells | bone marrow |
| 088b475a-17ae-56c3-a038-c9eb57da53af | 1M Immune Cells | bone marrow |
| 0bd29b3f-a5f4-5d70-ace4-4ecda6f00ebf | 1M Immune Cells | bone marrow |
| 0e60185d-ffad-5ceb-b822-7f40b57e1cba | 1M Immune Cells | bone marrow |
| 11dac858-6f2f-5e73-9824-428f677b0076 | 1M Immune Cells | bone marrow |
| 322f880b-aea5-53d4-b8ce-77e68ac6e3f3 | 1M Immune Cells | bone marrow |
| 3e977c03-cf2c-55a2-a012-3337c433dfc1 | 1M Immune Cells | bone marrow |
| 63d23827-d07c-58c1-8ec4-e9a8027721c6 | 1M Immune Cells | bone marrow |
| 6c9e3cf8-1d7d-5f38-a0be-ac588c6581ca | 1M Immune Cells | bone marrow |
| 6ce9adc1-4457-5a78-a50d-f4c14341656d | 1M Immune Cells | bone marrow |
| 83c5100c-716a-5cb2-a587-bff4860d97d4 | 1M Immune Cells | bone marrow |

| | | |
|---|---|---|
| 88945738-797b-55b9-a694-7e36077c2a1b | 1M Immune Cells | bone marrow |
| 8aed3195-e12e-5f30-beb3-658caf82d99f | 1M Immune Cells | bone marrow |
| 8be544ee-692e-5ee8-a4f0-4d674698f075 | 1M Immune Cells | bone marrow |
| 96604aad-3aa4-583d-9ab5-68359b7ed1b4 | 1M Immune Cells | bone marrow |
| a84816b0-8dc4-5e29-b817-dce68e6f323f | 1M Immune Cells | bone marrow |
| aa2af946-4a7e-57d8-b437-08ae17aeb61f | 1M Immune Cells | bone marrow |
| ad4d7505-9796-56c4-9dda-05005fb691e1 | 1M Immune Cells | bone marrow |
| b26f2473-70d9-540b-b134-8b18156006e7 | 1M Immune Cells | bone marrow |
| b4e31ea4-8e3d-5751-ab7a-6bb88e1ff2cc | 1M Immune Cells | bone marrow |
| be02548a-28ac-5240-b76f-849720700786 | 1M Immune Cells | bone marrow |
| c8af8721-b7d1-5976-a909-74d6461de682 | 1M Immune Cells | bone marrow |
| d45ba229-bf85-5133-a2d1-f20c95dbdacf | 1M Immune Cells | bone marrow |
| d838f3de-9c60-5e53-a900-466039b51feb | 1M Immune Cells | bone marrow |
| d89ecb27-06c2-5d40-8bf3-07433bae87e3 | 1M Immune Cells | bone marrow |
| da0fa4d4-ea92-588b-a15a-308d1c8604d5 | 1M Immune Cells | bone marrow |
| e76430e9-6ccd-52a5-b35c-71a1fe090ca1 | 1M Immune Cells | bone marrow |
| 84fc17e6-1eff-4e9c-8f31-28505ce881a9 | CrossTissueReferenceMap | skeletal muscle organ |
| 84fc17e6-1eff-4e9c-8f31-28505ce881a9 | CrossTissueReferenceMap | prostate gland |
| 84fc17e6-1eff-4e9c-8f31-28505ce881a9 | CrossTissueReferenceMap | lung |
| 84fc17e6-1eff-4e9c-8f31-28505ce881a9 | CrossTissueReferenceMap | heart |
| 84fc17e6-1eff-4e9c-8f31-28505ce881a9 | CrossTissueReferenceMap | esophagus |
| 616f5494-308a-528a-9313-774dc8e56ba6 | EpithelialDiversityHealthInflammation | colon |
| 4da82749-fac6-56cd-9f9c-b9c4e3e4434f | EpithelialDiversityHealthInflammation | colon |
| 272eb311-4e81-5444-b079-7a6a42e9d413 | EpithelialDiversityHealthInflammation | colon |
| 06f334bd-ac04-5e3e-b8e1-9e607fe8c3e7 | HeartSingleCellsAndNucleiSeq | heart |
| 230eaaee-beb7-584a-a3b3-fd1682c4785f | HeartSingleCellsAndNucleiSeq | heart |
| 3f8b4e5f-11b7-5362-bcbc-5e7142179b11 | HeartSingleCellsAndNucleiSeq | heart |
| 443281f9-a5c4-5b63-a704-c6daf0b02e0e | HeartSingleCellsAndNucleiSeq | heart |
| 4d07037f-6f59-54ea-bfe8-80b6297a6da6 | HeartSingleCellsAndNucleiSeq | heart |
| 5b5e6142-b5ea-5f2c-b485-e59ba7302a1d | HeartSingleCellsAndNucleiSeq | heart |
| a286a65b-a267-571e-a4eb-980e77ac8752 | HeartSingleCellsAndNucleiSeq | heart |
| a66f86fa-cbe4-5018-ab86-dccfed192679 | HeartSingleCellsAndNucleiSeq | heart |
| c3eef56d-96b7-5af4-a8b7-edbe9563306c | HeartSingleCellsAndNucleiSeq | heart |
| fb1b3d9b-8bc5-5e46-8801-4aac68c64e67 | HeartSingleCellsAndNucleiSeq | heart |
| 1f3d18c2-65c0-5d4e-8075-fc331b41b3ff | HeartSingleCellsAndNucleiSeq | heart |
| 23484bd9-5efc-537a-96e7-346bb7319ff8 | HeartSingleCellsAndNucleiSeq | heart |
| 2375387f-6786-5750-8559-f5dd73477348 | HeartSingleCellsAndNucleiSeq | heart |
| 2ca6b96e-a40f-52db-a925-4720223caaeb | HeartSingleCellsAndNucleiSeq | heart |
| 4abfa27d-449a-5ac7-9ca1-ecb9dba0a02a | HeartSingleCellsAndNucleiSeq | heart |
| 6f0b7799-7db3-5d5a-bf0a-ade76c996a5a | HeartSingleCellsAndNucleiSeq | heart |
| 7821e415-43fa-55b8-b656-b8c80059168c | HeartSingleCellsAndNucleiSeq | heart |
| 78715aed-6e65-5a5e-ab04-4a6147d08c11 | HeartSingleCellsAndNucleiSeq | heart |
| 9527526f-33ac-5a95-adfe-b39136e3b139 | HeartSingleCellsAndNucleiSeq | heart |
| 9582cba1-217b-564b-87fc-8557d0bccd52 | HeartSingleCellsAndNucleiSeq | heart |
| ae7c7470-6ae6-5486-9ee6-bfd4433b540e | HeartSingleCellsAndNucleiSeq | heart |
| 0110b875-60d3-582a-928e-1f58d1a70de5 | HeartSingleCellsAndNucleiSeq | heart |
| 0aa51b10-b2ec-512c-9188-452c34f9f788 | HeartSingleCellsAndNucleiSeq | heart |

| | | |
|---|---|---|
| 195b7cfd-172a-5eb0-adee-d925a296bfbf | HeartSingleCellsAndNucleiSeq | heart |
| 3515d736-9d29-58bb-9eeb-616cf8f50d46 | HeartSingleCellsAndNucleiSeq | heart |
| 3d456927-14b2-5983-b7b6-eb5c2d43deaf | HeartSingleCellsAndNucleiSeq | heart |
| 4c3184b5-05ef-5989-9fba-b801fcceeab6 | HeartSingleCellsAndNucleiSeq | heart |
| 4eba3717-518a-5af8-bd68-53f67c32ccb6 | HeartSingleCellsAndNucleiSeq | heart |
| 510daec2-fa5e-55eb-867b-cfd4b3cb367e | HeartSingleCellsAndNucleiSeq | heart |
| 6942ecb2-d6d9-5451-bc03-1e2f93257212 | HeartSingleCellsAndNucleiSeq | heart |
| 8d12547f-0fc5-529a-b426-756b9c53e70e | HeartSingleCellsAndNucleiSeq | heart |
| b7cbed24-5441-5105-8244-e40bb370ae16 | HeartSingleCellsAndNucleiSeq | heart |
| c9cb3405-ca4e-5fd4-93fb-e6a7a8cda833 | HeartSingleCellsAndNucleiSeq | heart |
| cbed9f1f-34b6-515d-8080-b5c019097fc0 | HeartSingleCellsAndNucleiSeq | heart |
| ccb97ac0-d16d-50f9-ad73-5a6e5b4647a2 | HeartSingleCellsAndNucleiSeq | heart |
| d5797a79-319d-5836-9405-644d869cde2e | HeartSingleCellsAndNucleiSeq | heart |
| d6f91a43-0e2e-59ca-a2f2-3960038073fb | HeartSingleCellsAndNucleiSeq | heart |
| d85b75aa-f12f-5a73-8b4d-09c928a2bf94 | HeartSingleCellsAndNucleiSeq | heart |
| dae85cda-ace0-558a-ae9c-bf2fc2de8679 | HeartSingleCellsAndNucleiSeq | heart |
| e8237984-25c4-5514-a2ec-a166acb380c5 | HeartSingleCellsAndNucleiSeq | heart |
| f2c87df8-1dd7-53c2-aab1-ca7d7a111d58 | HeartSingleCellsAndNucleiSeq | heart |
| c4ed9481-ddfc-5869-aa0a-20e8bb94d744 | HumanBrainSubstantiaNigra | brain |
| 734f9853-f39b-543f-bf9c-342d9431a4ac | HumanBrainSubstantiaNigra | brain |
| c9b45b64-10c7-56bc-a2c9-a8ef07acac17 | HumanBrainSubstantiaNigra | brain |
| 89771a00-45d9-5ccb-b3b8-e1f316753905 | HumanBrainSubstantiaNigra | brain |
| ce16e9e3-a705-5b8a-86b0-ae882679ab80 | HumanBrainSubstantiaNigra | brain |
| 8cd6b2b3-32b4-5a1f-8562-61ee76631c62 | HumanBrainSubstantiaNigra | brain |
| dbe3a518-b452-5b0d-b856-aa958586ce7d | HumanBrainSubstantiaNigra | brain |
| aacc0f02-9d8d-5466-b5ce-c567afd21f9a | HumanBrainSubstantiaNigra | brain |
| dc6af900-2540-5bda-8113-ccb8d22891cb | HumanLiverImmuneCells_GSE125188 | spleen |
| d660be85-872c-5092-b26e-2e6fb48b0fa9 | HumanLiverImmuneCells_GSE125188 | spleen |
| c14a27e0-f7f9-5334-9c78-459a6d398ecb | HumanLiverImmuneCells_GSE125188 | spleen |
| 15b56725-e92f-5fe6-8fb8-a3360ea3c94c | HumanLiverImmuneCells_GSE125188 | liver |
| f0b69f95-2145-5e78-8f45-48d503b92792 | HumanLiverImmuneCells_GSE125188 | liver |
| 00ef76b5-455c-5a62-8bfe-9ef308516b03 | HumanLiverImmuneCells_GSE125188 | liver |
| 8567b540-bc7b-5e6a-b0f2-1a4abe3f2545 | HumanLiverImmuneCells_GSE125188 | blood |
| b6165a2f-29ef-5f64-b9fa-b7b92b63c543 | HumanLiverImmuneCells_GSE125188 | blood |
| 87613813-14fe-5a44-9c51-b07fb3eed9b6 | HumanThymicDevelopment | thymus |
| 3fc18055-e6ec-5cf6-af36-8ea9be272ff5 | HumanThymicDevelopment | thymus |
| e10ed38b-dbef-55c4-af7b-83ecf5f890c4 | HumanThymicDevelopment | thymus |
| 625add28-2566-57e6-be48-094abaa768c8 | HumanThymicDevelopment | thymus |
| 9a9b2c08-7397-5072-8d33-53b8adcc3f03 | HumanThymicDevelopment | thymus |
| 92a5e8c5-85c1-5be8-95fe-e3a56c594a03 | HumanThymicDevelopment | thymus |
| 12f18411-99dd-5871-b142-10fc68de8a58 | HumanThymicDevelopment | thymus |
| a5c9ec18-c173-52b4-8046-e7cf147790f1 | HumanThymicDevelopment | thymus |
| 71b3c1e4-2e09-5d78-bea7-09d87416cb66 | KidneySingleCellAtlas | kidney |
| 55f6e84c-e01e-5174-9237-973ea9022041 | KidneySingleCellAtlas | kidney |
| 9c052ef5-b1f6-5520-9d3b-f1b3a6a49505 | KidneySingleCellAtlas | kidney |
| 91520010-7082-58af-9feb-cefbe272f550 | KidneySingleCellAtlas | kidney |
| 2b4ed392-328c-59ae-8102-8ba9e7ee7594 | KidneySingleCellAtlas | kidney |

| | | |
|---|---|---|
| 1fe6ce43-231a-57c8-b98a-4b68b6c0ba3d | KidneySingleCellAtlas | kidney |
| f4770d58-6e1e-57c8-907f-cc4234e5bc1a | KidneySingleCellAtlas | kidney |
| 20ba2d6c-7eaa-58de-a9b0-dbd8c63de8ad | KidneySingleCellAtlas | kidney |
| 6c2bab8f-dec0-51f6-87bc-479e1bf3e429 | ProstateCellAtlas | prostate gland |
| 43357eae-eb10-56dc-a199-b88a0c3e5c39 | ProstateCellAtlas | prostate gland |
| 43a2301f-e975-5872-b057-63ca8262987d | ProstateCellAtlas | prostate gland |
| 0f0fce48-b6b9-54a1-af11-caf7289cf33b | ProstateCellAtlas | prostate gland |
| c4bc6970-df83-58c4-81b3-580e75104afb | ProstateCellAtlas | prostate gland |
| e084a288-f027-5283-8337-7df0411d7122 | ProstateCellAtlas | prostate gland |
| 225d2456-7954-5e16-a6a7-87edf3de8106 | ProstateCellAtlas | prostate gland |
| b503b3d0-a2bf-5852-8e1e-452ba08ee3ec | ProstateCellAtlas | prostate gland |
| 96b6f7ee-b736-52b2-af30-b944a560a82b | ProstateCellAtlas | prostate gland |
| 5c72120d-0609-5007-8812-cef855a0a35e | ProstateCellAtlas | prostate gland |
| b79c4b81-8e48-5dee-9af0-980e6c8fab19 | SkeletalMuscleHumanMouse | skeletal muscle organ |
| e2d27a07-706e-563e-89cd-b1e1cf90944f | SkeletalMuscleHumanMouse | skeletal muscle organ |
| be109b77-46ce-58c6-b065-9ae603436100 | SkeletalMuscleHumanMouse | skeletal muscle organ |
| 0b17d08b-bc94-5976-b519-45e06fb8671f | SkeletalMuscleHumanMouse | skeletal muscle organ |
| 09683288-b9be-5cc1-9414-f3e138fe355d | TCellsHumanCentralNervousSystem | brain |
| 28fd2e91-5a0c-565c-a0ba-ce944309353e | TCellsHumanCentralNervousSystem | brain |
| 4cf3f070-a0b4-55e8-b9fe-7e62ef9eeceb | WongAdultRetina | eye |
| a0814d48-4082-5bef-9c9a-1584a5133b87 | WongAdultRetina | eye |
| d535b0a4-e3be-59b2-afef-c37567f455c5 | WongAdultRetina | eye |
| 7eb74d9f-8346-5420-b7e4-b486f99451a8 | WongAdultRetina | eye |

| Sex | Age | Condition |
|---|---|---|
| female | 36 year | healthy donor |
| female | 32 year | healthy donor |
| female | 36 year | healthy donor |
| female | 32 year | healthy donor |
| female | 52 year | healthy donor |
| male | 29 year | healthy donor |
| female | 36 year | healthy donor |
| female | 52 year | healthy donor |
| female | 52 year | healthy donor |
| male | 50 year | healthy donor |
| male | 29 year | healthy donor |
| female | 36 year | healthy donor |
| male | 29 year | healthy donor |
| female | 36 year | healthy donor |
| male | 29 year | healthy donor |
| female | 32 year | healthy donor |
| male | 39 year | healthy donor |
| male | 29 year | healthy donor |
| male | 29 year | healthy donor |
| male | 39 year | healthy donor |
| female | 32 year | healthy donor |
| female | 26 year | healthy donor |
| male | 29 year | healthy donor |
| male | 29 year | healthy donor |
| male | 50 year | healthy donor |
| female | 52 year | healthy donor |
| male | 29 year | healthy donor |
| male | 29 year | healthy donor |
| male | 29 year | healthy donor |
| female | 32 year | healthy donor |
| female | 52 year | healthy donor |
| male | 50 year | healthy donor |
| female | 26 year | healthy donor |
| female | 52 year | healthy donor |
| male | 50 year | healthy donor |
| male | 39 year | healthy donor |
| male | 39 year | healthy donor |
| male | 39 year | healthy donor |
| male | 29 year | healthy donor |
| female | 26 year | healthy donor |
| male | 50 year | healthy donor |
| male | 29 year | healthy donor |
| male | 50 year | healthy donor |
| male | 39 year | healthy donor |
| male | 39 year | healthy donor |

| | | |
|---|---|---|
| female | 26 year | healthy donor |
| male | 29 year | healthy donor |
| female | 52 year | healthy donor |
| female | 26 year | healthy donor |
| male | 50 year | healthy donor |
| female | 36 year | healthy donor |
| female | 32 year | healthy donor |
| female | 32 year | healthy donor |
| female | 52 year | healthy donor |
| female | 26 year | healthy donor |
| female | 26 year | healthy donor |
| male | 29 year | healthy donor |
| female | 36 year | healthy donor |
| male | 39 year | healthy donor |
| male | 29 year | healthy donor |
| female | 36 year | healthy donor |
| NA | NA | healthy donor |
| NA | NA | healthy donor |
| NA | NA | healthy donor |
| NA | NA | healthy donor |
| NA | NA | healthy donor |
| NA | NA | healthy donor |
| NA | NA | healthy donor |
| NA | NA | healthy donor |
| male | 60-65 year | healthy donor |
| female | 60-65 year | healthy donor |
| male | 60-65 year | healthy donor |
| male | 65-70 year | healthy donor |
| male | 65-70 year | healthy donor |
| male | 60-65 year | healthy donor |
| male | 55-60 year | healthy donor |
| male | 60-65 year | healthy donor |
| male | 55-60 year | healthy donor |
| female | 60-65 year | healthy donor |
| female | 60-65 year | healthy donor |
| male | 60-65 year | healthy donor |
| male | 60-65 year | healthy donor |
| female | 60-65 year | healthy donor |
| female | 60-65 year | healthy donor |
| male | 55-60 year | healthy donor |
| female | 60-65 year | healthy donor |
| male | 65-70 year | healthy donor |
| female | 60-65 year | healthy donor |
| female | 65-70 year | healthy donor |
| male | 65-70 year | healthy donor |
| female | 65-70 year | healthy donor |
| female | 65-70 year | healthy donor |

| | | |
|---|---|---|
| male | 60-65 year | healthy donor |
| male | 60-65 year | healthy donor |
| female | 65-70 year | healthy donor |
| male | 55-60 year | healthy donor |
| male | 60-65 year | healthy donor |
| male | 60-65 year | healthy donor |
| female | 65-70 year | healthy donor |
| male | 65-70 year | healthy donor |
| male | 65-70 year | healthy donor |
| male | 65-70 year | healthy donor |
| female | 60-65 year | healthy donor |
| male | 60-65 year | healthy donor |
| male | 65-70 year | healthy donor |
| female | 65-70 year | healthy donor |
| male | 65-70 year | healthy donor |
| male | 55-60 year | healthy donor |
| female | 60-65 year | healthy donor |
| female | 60-65 year | healthy donor |
| NA | NA | healthy donor |
| NA | NA | healthy donor |
| NA | NA | healthy donor |
| NA | NA | healthy donor |
| NA | NA | healthy donor |
| NA | NA | healthy donor |
| NA | NA | healthy donor |
| NA | NA | healthy donor |
| male | 27 year | healthy donor access to liver transplantation |
| male | 41 year | healthy donor access to liver transplantation |
| male | 56 year | healthy donor access to liver transplantation |
| male | 27 year | healthy donor access to liver transplantation |
| male | 41 year | healthy donor access to liver transplantation |
| male | 56 year | healthy donor access to liver transplantation |
| male | 27 year | healthy donor access to liver transplantation |
| male | 56 year | healthy donor access to liver transplantation |
| female | 20-25 year | healthy donor |
| female | 20-25 year | healthy donor |
| female | 20-25 year | healthy donor |
| female | 20-25 year | healthy donor |
| female | 20-25 year | healthy donor |
| female | 20-25 year | healthy donor |
| female | 20-25 year | healthy donor |
| female | 20-25 year | healthy donor |
| male | 72 year | healthy donor |
| female | 53 year | healthy donor |
| female | 53 year | healthy donor |
| male | 72 year | healthy donor |
| male | 72 year | healthy donor |

| | | |
|---|---|---|
| female | 53 year | healthy donor |
| female | 53 year | healthy donor |
| female | 53 year | healthy donor |
| male | 31.0 year | healthy donor |
| male | 25.0 year | healthy donor |
| male | 31.0 year | healthy donor |
| male | 25.0 year | healthy donor |
| male | 25.0 year | healthy donor |
| male | 25.0 year | healthy donor |
| male | 29.0 year | healthy donor |
| male | 25.0 year | healthy donor |
| male | 29.0 year | healthy donor |
| male | 25.0 year | healthy donor |
| male | 26 year | healthy donor |
| male | 26 year | healthy donor |
| male | 26 year | healthy donor |
| male | 26 year | healthy donor |
| male | 41.0 year | healthy donor |
| male | 41.0 year | healthy donor |
| male | 42 year | healthy donor |
| female | 53 year | healthy donor |
| female | 53 year | healthy donor |
| male | 42 year | healthy donor |
